# Neural Mediators of Altered Perceptual Choice and Confidence Using Social Information

**DOI:** 10.1101/516963

**Authors:** Tiasha Saha Roy, Bapun Giri, Arpita Saha Chowdhury, Satyaki Mazumder, Koel Das

**Affiliations:** Department of Mathematics and Statistics, Indian Institute of Science Education and Research Kolkata, Mohanpur, Nadia-741246, India; Department of Psychology, University of Wisconsin-Milwaukee, PO Box 413, Milwaukee, WI 53201, USA; Department of Anesthesiology, University of Michigan, Ann Arbor, Michigan 48109, USA

**Keywords:** Perceptual decision making, Social influence, Computational modeling, Gamma mixture model, Multivariate pattern classification

## Abstract

Understanding how individuals utilize social information while making perceptual decisions and how it affects their decision confidence is crucial in a society. Till date, very little is known about perceptual decision making in humans under the influence of social cues and the associated neural mediators. The present study provides empirical evidence of how individuals get manipulated by social cues while performing a face/car identification task. Subjects were significantly influenced by what they perceived as decisions of other subjects while the cues in reality were manipulated independently from the stimulus. Subjects in general tend to increase their decision confidence when their individual decision and social cues coincide, while their confidence decreases when cues conflict with their individual judgments often leading to reversal of decision. Using a novel statistical model, it was possible to rank subjects based on their propensity to be influenced by social cues. This was subsequently corroborated by analysis of their neural data. Neural time series analysis revealed no significant difference in decision making using social cues in the early stages unlike neural expectation studies with predictive cues. Multivariate pattern analysis of neural data alludes to a potential role of frontal cortex in the later stages of visual processing which appeared to code the effect of social cues on perceptual decision making. Specifically medial frontal cortex seems to play a role in facilitating perceptual decision preceded by conflicting cues.

## Introduction

In todays information-satiated society, perceptual decision and subsequent action is greatly influenced by social information. Modern human society is increasingly organized around collective opinions reflected in peoples increased use of web ratings for daily choices about consumer products, lodging, food and entertainment [1]. Opinions and choice can easily propagate through social networks [2, 3] in this digitized world and even political opinions can be manipulated using social transmission [4]. Human tendency to conform to social influence has been explored systematically in classic studies by Solomon Asch [5, 6] and others ([7–15] and see [16–18] for reviews). Reliance on others opinion is not unique to humans and different species of animal depend on collective opinion to decide life-critical perceptual tasks like foraging for food, placement of their nests and navigation [19–21] and evolve optimal decision strategies accordingly. Beneficial effect of group decision can be traced as early as 1907 when Francis Galton analyzed the opinions of 787 people about the weight of an ox and found that combining their numerical assessments resulted in a median estimate that was remarkably close to the true weight of the ox [22]. In recent times, this idea has been popularly referred to as the wisdom of the crowds [23]. However, effect of social cues in the form of collective decision on individual percept and the underlying neural mechanism remains largely unexplored [12, 18].

Neural expectation studies over the last decade have demonstrated that predictive cues typically lead to changes in early sensory processing [24–34] but recent research has contradicted this claim [35, 36]. We sought to examine whether social information produces similar early top down changes in sensory cortex. We propose to manipulate the individual choice and decision confidence of humans performing a perceptual task by presenting visual cues which the subjects presumed to be collective opinion of other well performing participants. The social cues can be concurring, conflicting or neutral to the individual perceptual decision of the subjects. Using a novel statistical model, we studied the effect of the three types of social cues on their individual choice. We also analyzed the neural signals to explore the neural mediators producing the change in their individual choice upon presenting social cues. Finally we performed a source reconstruction of the neural signals to elucidate the role played by specific spatio-temporal areas under the influence of social cues. Specifically we explored the following questions:

Can we manipulate individual perceptual decision upon presenting social information cues when the social decision differs from the individual choice? Does this reversal of opinion depend upon how confident the subject was in his/her choice without the social information?

Can the individual decision confidence be augmented when the social cues concur with the individual choice?

Can we identify the flip-floppers based on computational modeling of their behavioural data and corroborate using neural data?

Can we explore the neural mediators that contribute to the change in individual percept post social information?

Using a face/car discrimination task, we show that it is possible to manipulate individual choice post presentation of social cues in the guise of others decision. Although the social cues were randomly generated and independent from the stimulus, it was possible to alter the individual percept as subjects presumed the social cues as concurring, conflicting or neutral. Irrespective of the order in which they viewed the images with or without social cues, most subjects were affected by the social cues in a systematic manner. The distribution of the decision confidence under such set up was found to be bi-modal and skewed with one mode guided by social cues and the other influenced by their own decision. The tendency to adhere to their own decision depends on the confidence level of the subject and is reflected in the skewness of the data distribution. Hence using a Gaussian model to explore the data, which is the usual practice [37], might not capture the complexities of data completely. We propose a novel model using a mixture of shifted gamma and negative gamma distribution which successfully captures the effect of social cues on individual choice. To the best of our knowledge, this is the first work using a mixture of variants of gamma distributions which captures the bi-modal nature as well as the skewness (whether high or low) of this kind of data. We compare our proposed model with bi-modal Gaussian and demonstrate the superiority of our model convincingly. Based on the behavioural model, it was possible to objectively identify subjects most prone to change their decisions upon presenting others’ opinion. Subsequent multivariate pattern analysis (MVPA) of neural data substantiated the above finding. Neural analysis also elucidated existence of a late component that seem to code the effect of this social information on individual perceptual decision. Source analysis of neural data revealed a role of frontal cortex in coding perceptual decision using social information. Our analysis alludes to the role of medial frontal cortex in coding information when conflicting social decisions are provided as cues.

## Materials and Methods

### Ethics Statement

This study was carried out in accordance with the recommendations of ‘Institute Ethics Committee’ at Indian Institute of Science Education and Research Kolkata, India with written informed consent from all subjects. All subjects gave written informed consent in accordance with the Declaration of Helsinki. The protocol was approved by the ‘Institute Ethics Committee’.

### Stimuli and display

The data set consisted of 290×290 pixel 8-bit gray-scale images of 12 cars and 12 faces with equal number of frontal views and side views. Face images were taken from the Max Planck Institute for Biological Cybernetics face database [38]. All stimuli were filtered to attain a common frequency power spectrum. Noise was generated by filtering white Gaussian noise (std of 3.53 cd/m2) by the average power spectrum. Noise was added to the base stimuli to generate a set of 250 images (125 face, 125 car). Contrast energy of all 250 images was matched at 0.3367 *deg* ^*2*^. The participants were at a distance of 125 cm from the display with a mean luminance of 25 cd/ *m*^2^. Images subtended a visual angle of 4.57 degree.

### Participants and Experiment

Twenty na’ive participants (ages: 22-28 mean: 25.85 std: 2.39) participated in the study which consisted of 1000 trials split into 40 successive sessions. Three subjects were not considered in the analysis due to high degree of noise present in the neural data. All participants had normal or corrected-to-normal vision and disclosed no history of neurological problems. The participants performed a face/car discrimination task and reported their decision using a 10-point confidence rating. Participants perceptually categorized briefly (50 ms) presented images of cars (C) and faces (F) embedded in filtered noise. The participants began by fixating on a central cross and clicking anywhere on the screen. After a delay of 50 ms, a cue was presented for 100 ms followed by a variable delay of 500-800 ms. The stimulus was presented for 50 ms followed by delay of 700 ms after which the response screen appeared. The participants reported their decision using the confidence rating with a rating of 1 indicating complete confidence that the stimuli was a face and a rating of 10 indicating that it was a car with complete confidence. The participants reported their confidence rating on a grey-scaled colorwheel in the response screen to avoid any motor bias (Fig. 1A). There were four types of cues, FF, *CC*, FC, CF, representing decisions of two independent well performing participants who had previously completed the study. Cues were systematically manipulated such that equal number of images (250 per condition) have FF cues, FC/CF cues and CC cues. There were also additional 250 images without cues. Thus each participant saw one stimuli four times preceded by FF cue, FC/CF cue, CC cue and no cue in the course of the experiment in random order and the responses were recorded. Participants were naïve to the purpose of the study and in subsequent questionnaire after the study failed to realize that the cues were not decision cues and were in fact synthetic cues generated randomly.

**Figure 1:**
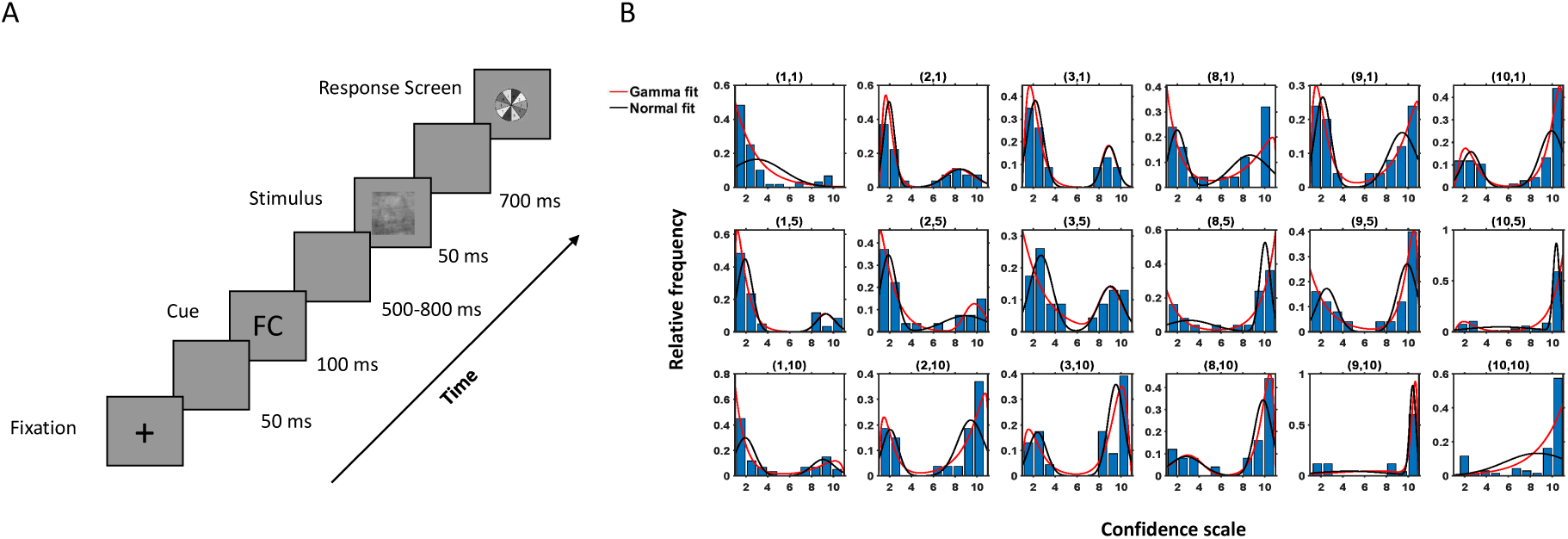
Experimental protocol and behavioural response. (**A**) Experimental Paradigm. (**B**) Histogram of the observed data and fitted density of the proposed model (red) and Gaussian mixture model (black) for a subject for different combinations of *k*_1_, *k*_2_ (denoted on top of each case, e.g. (1,10) implies subject data and fitted model for the images when individual choice was 1 denoting face with highest confidence and social cue was CC). Here x-axis denotes the confidence scale and y-axis denotes the relative frequencies of the subject’s choices for a particular combination of *k*_1_, *k*_2_.

EEG activity was recorded using a 64 channel active shielded electrodes mounted in an EEG cap following the international 10/20 system. EEG signals were recorded using 2 linked Nexus-32 bioamplifliers at a sampling rate of 512 Hz., band-pass filtered (0.01 − 40 Hz.) and then referenced using average referencing. Trials with ocular artifacts (blinks and eye movements) were detected using bipolar electro-occulograms (EOG) with amplitude exceeding ±100 mV or visual inspection and not included in the analysis.

### Behavioural model

We propose a statistical model to explore the effect of social cues on perceptual decision making. In the experiment, for every face/car stimuli, subject responses corresponding to the three types of social cues (FF, FC/CF and CC) along with a response to the same stimuli with no-cues were recorded. The response to the no-cue image was taken as the individual decision of the subject, *k*_1_ ∈ {1, 2, … 10}, for that image. Further, we define a social cue variable *k*_2_ as

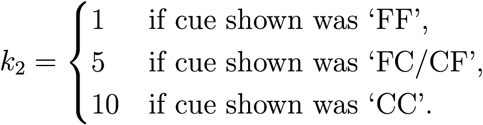

All the images in which the individual decision of the subject was *k*_1_, were considered and the distribution of the decisions on the same images under the influence of each type of social cue was studied. Hence the data comprised of the decisions of a particular subject for every (*k*_1_,*k*_2_) pair. In most cases, the data distributions were bimodal in nature having positive and/or negative skew, as seen in Fig. 1B. Hence a two-component mixture model based on variants of the gamma distribution was proposed to explain the decisions taken by the subject under the influence of a social cue. The data was made continuous by using jittering (addition of uniform random noise, [39]) to provide flexibility in modeling.

Let **X**_*i*_(*k*_**1**_, *k*_2_) contain the decisions taken by the *ith* subject on all images, where his/her individual decision was *k*_1_ and cue shown was *k*_2_. We consider the elements of **X**_*i*_(k_**1**_, *k*_2_) as i.i.d. observations from a distribution. To propose the statistical model depending on the choices of (*k*_1_, k_2_) we first introduce some terminology and notation. The probability densities of *shifted gamma* and *negative gamma* distributions are given respectively as

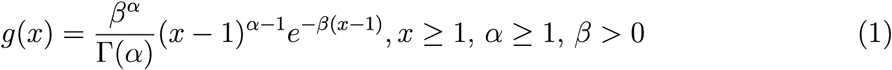

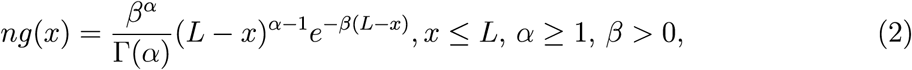

where *α* and *β* are the shape and scale parameters respectively and *L* is a known constant.

Based on the equations (1) and (2) the following models are proposed depending on the choices of (*k*_1_, *k*_2_). If *k*_1_ ∈ {1, 2, …, 5} and *k*_2_ ∈ {1, 5} we take our model as

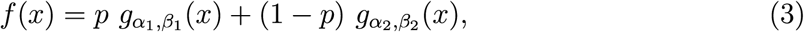

a mixture of two shifted gamma distributions. When *k*_1_ ∈ {6, 7 …, 10} and *k*_2_ = 10 the proposed model is

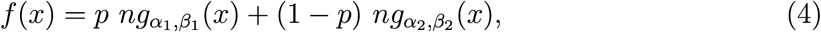

a mixture of two negative gamma distributions. Finally if either *k*_1_ ∈ {1, 2, …, 5} and *k*_2_ = 10 or *k*_1_ ∈ {6, 7 …, 10} and *k*_2_ ∈ {1, 5} our suggested model is

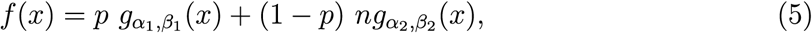

a mixture of a shifted gamma and a negative gamma distribution, where 0 ≤ *p* ≤ 1 is the mixing parameter.

#### Parameter space of the model

We have taken the restricted parameter space for the shape parameter (*α*) in both the distributions (equations (1) and (2)) so that mode of the distribution is defined and either that is more than or equal to 1 (for shifted gamma case) or that is less than or equal to *L* (for negative gamma case). In our case, we consider *L* to be 11. In particular for both shifted-gamma and negative-gamma distributions,

- the shape parameter *α* ∈ [1, ∞) and
- the scale parameter *β* ∈ (0, ∞).

#### Estimation of the model parameters

Next, for the purpose of estimation of parameters of our proposed model and further inference, only those data are considered which have more than 10 observations. Note that the parameter estimates depend on *i* as well as (*k*_1_, *k*_2_), that is to say, for every individual *i*, the parameter estimates may vary for different choices of (*k*_1_, *k*_2_). Similarly for a given (*k*_1_, *k*_2_), parameter estimates of the proposed model may vary from individual to individual. We estimate the model parameters by maximum likelihood estimation procedure [40]. Since the proposed models are mixture densities, so to calculate the maximum likelihood estimates (MLE) we invoke the technique of EM algorithm [40]. However, since closed form solution for estimates of shape parameters do not exist, we apply Newton Raphson numerical technique [41] within each M-step of the EM-algorithm (see Supporting Information for detailed calculation).

#### Goodness of fit

To understand how well our model fits the observed data, Kolmogorov-Smirnov (KS) test statistic [42], based on the maximum absolute differences between the hypothesized cumulative distribution function (cdf) and empirical cumulative distribution function (ecdf) was used. For each subject *i*, there were *N*_*i*_ models to be tested simultaneously and therefore arose the case of multiple testing. To control the family wise error rate, arising due to multiple hypotheses testing per subject, we used the Holm-Bonferroni method [43] with a family-wise error rate (FWER) of 0.05.

#### Model prediction

We use 10-fold cross validation procedure to study the predictive performance of the proposed model. Since our data was bimodal in nature, it would not have been meaningful to judge this performance on the basis of a single predictive interval. To address this issue, we apply the following concept of highest probability density region (HPDR) [44] which broadly computes the smallest region that contains most of the probability.

##### Definition

Let *f(x)* be the probability density function of a random variable *X*. Then the 100(1-α)% HPDR is defined as the subset R(*f*_*α*_) of real numbers, ℝ, such that

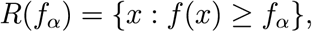

where *f*_*α*_ is the largest constant with *P(X* ∈ *R*(*f*_*α*_)) 1 − α.

In each fold, model was trained on the training set and the 95% HPDR was computed. It was checked whether the validation set falls in the estimated HPDR and the process was repeated for each cross-validation fold.

#### Model comparison

We compared the performance of our proposed model with the 2-component Gaussian mixture model using likelihood ratio test [40]. Data was divided into 10 test sets using 10-fold cross validation and for each set the likelihood was estimated from each of the two models. Finally, the medians of the likelihood ratios across the folds were computed for each of the models for the purpose of comparison.

### Behavioural data processing

Guided by the proposed model the behaviour of the individuals was analyzed based on the following measures.

#### Distance metric computation using the model

To quantify the overall shift in decisions from the subjects’ individual choice, the following distance was used

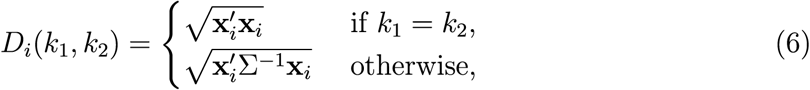

where **X**_*i*_ = (*k*_1_ - **m**_**1**_ (*i*), *k*_1_ - **m**_**2**_ (*i*))′, **m**_**1**_ and **m**_**2**_ being the vectors containing the two modes of the 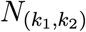 subjects and 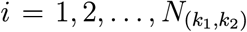. Here 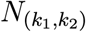 denotes the number of subjects available corresponding to (*k*_1_, *k*_2_) and Σ is the estimated variance covariance matrix of estimates of the modes for a particular choice of (*k*_1_, *k*_2_), given by

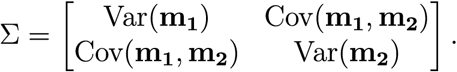

#### Social Bias Score

Using the cumulative distribution functions of shifted-gamma and negative gamma distributions (as calculated in SI) and equations (3)–(5) the proportion of decisions between *k*_1_ and *k*_2_ in presence of social cues was estimated. The average proportion of decisions (*P*_*i*_) per subject across the (*k*_1_, *k*_2_) pairs that have been reported in tables S5–S8 is considered. We rank the subjects based on social bias score, defined as

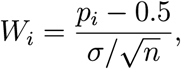

for *i* ∈ {1, 2, …, 17}\{2, 3} with *σ* denoting the sample standard deviation of the proportions *P*_*i*_. Only those subjects were considered for further analysis whose *W*_i_ exceeds 1.96, indicating that the corresponding proportions are significantly more than chance.

### Neural data processing

The preprocessed EEG signals were time-locked to stimulus onset and included a 200 ms pre-stimulus baseline and 500 ms post-stimulus interval.

#### Multivariate pattern analysis of EEG

Univariate EEG analysis had been used traditionally to explore the relationship between behavorial performance and neural activity in specific cognitive tasks. But the univariate analysis techniques fail to fully utilize the spatio-temporal nature of the multivariate neural data. Multivariate pattern analysis techniques provide a way of integrating the spatial and temporal information present in the data by fusing the neural information into a single decision variable which can be used in single trial analysis. A comparison of univariate analysis and multivariate analysis using a similar cognitive task has been shown in [45]. Successful use of MVPA has been demonstrated in numerous studies using EEG and fMRI [46–48]. In the current study, MVPA was used to extract meaningful information from the multi dimensional EEG data. Since the neural data is high dimensional and suffers from small sample size problem [49], a recently proposed principal component analysis (PCA) based non-linear feature extraction technique -’Classwise Principal Component Analysis –’ (CPCA) [49] is used. CPCA been used previously to efficiently reduce the dimensionality of the EEG signals and extract informative features [45, 50–54]. The main goal of CPCA is to identify and discard non-informative subspace in data by applying principal component based analysis to each class. The classification is then carried out in the residual space, in which small sample size conditions and the curse of dimensionality no longer hold. Linear Bayesian Classier was then used for computing the choice probability for single trial EEG data for each subject. Pattern analysis was performed using 10-fold cross validation. The original data was partitioned into 10 equal size subsamples. Of the 10 subsamples, a single subsample was retained as the test data, and the remaining 9 subsamples were used in training the classifier. The performance of the classifier is captured by the receiver operating characteristics (ROC) curve which plots the true positive rate vs. false positive rate at different classification thresholds. The area beneath this ROC curve (AUC) is often used as a measure to determine the overall accuracy of the classifier [55]. We utilize the well-known approach of calculating the area under the ROC by finding the MannWhitney U-statistic for the two-sample problem [56]. All classification analyses were carried out for individual participants and the average AUC performance was reported in the results.

#### Source Reconstruction

To identify underlying neuronal sources responsible for generating differences in the ERPs corresponding to the face and car trials under the influence of cues, source reconstruction was performed using sLORETA software([57], http://www.uzh.ch/keyinst/loreta). sLORETA (standardized low resolution brain electromagnetic tomography) is based on standardization of the minimum norm inverse solution which considers the variation of actual sources and the variation due to noisy measurement (if any) as well [5 7]. As a result, it does not have any localization bias even in the presence of measurement and biological noise. The head model for the inverse solution uses the electric potential lead field calculated using the boundary element method [58] on the MNI152 template [59]. The cortical grey matter is partitioned into 6239 voxels at 5 mm spatial resolution. sLORETA images represent the standardized electric activity at each voxel in Montreal Ne urological Institute (MNI) space as the exact magnitude of the estimated current density. Anatomical labels are reported using an appropriate correction from MNI to Talairach space [60] using Talairach Daemon [61]. For further details on sLORETA refer to http://www.uzh.ch/keyinst/NewLORETA/Methods/MethodsSloreta. The source activity was estimated from the face-car difference wave post stimulus onset.

#### Statistical Analysis of Sources

Differences in the distribution of the sources between concurring and conflict in g trials were calculated using statistical non-parametric mapping (SnPM) [62]. This method relies on the randomization of the absolute maximum-statistic over all channels. The randomization provides an estimator for the empirical distribution under the null hypothesis (“no difference between the sources of concurring and conflicting trials”). The advantage of this method is that it does not depend on any distributional form, in particular Gaussianity, and simultaneously takes care of multiple comparisons. 5000 random samples were generated while implementing the SnPM technique. Differences between the two conditions (concurring and conflicting) were assessed at the global level and the brain areas showing the largest differences have been reported.

## Results

### Behavioural Results

The decisions taken by the subjects under the influence of a social cue was modeled as a 2-component mixture model based on the shifted gamma and negative gamma distribution (see equations (3) – (5)). To verify that the proposed model fits the observed behaviour data well, the Kolmogorov-Smirnov (KS) test [42] was used. The proposed model captures the data correctly in most cases (see Table S1). Fig. 1B depicts the histogram of all (*k*_1,_ *k*_2)_ pairing and the fitted density of our model for one subject. Table S1 contains the p-values corresponding to the cases where the model is rejected. In over 96% of the cases, the hypothesized model was accepted, thus proving the efficacy of the model.

To estimate the predictive performance of the proposed model and prevent possible over-fitting, the highest probability density region (HPDR) of the fitted model was computed based on the training data and checked whether the test data falls in the calculated HPDR. Table S2 showing mean prediction error rates across subjects, demonstrate that the cross-validation error rate never exceeds 5% for any fold thus validating the excellent performance of the model in terms of prediction and nullifying the chance of over-fitting. Fig. 2A shows a fitted density function and the corresponding HPDR calculated from the training data of a particular validation fold of one subject. The test data as seen from the figure falls convincingly inside the indicated HPDR.

**Figure 2:**
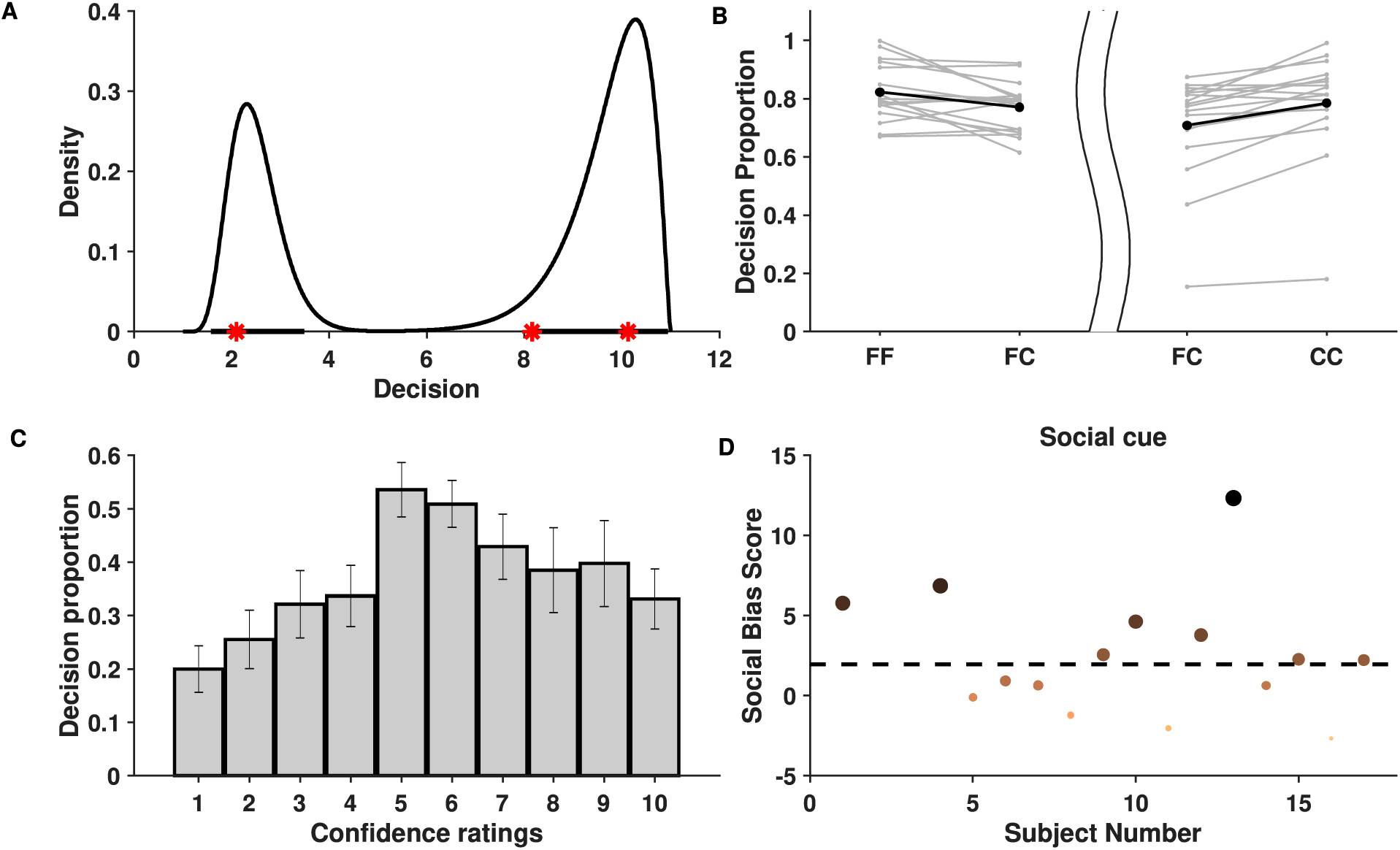
Behavioural Data Analysis. (**A**) Estimated probability density function based on the training data set shown for one subject when *k*_1_ = 3 and *k*_2_ = 10. Bold lines in x-axis represent the 95% HPDR and red stars represent the test observations for a subject. The test observations fall within the HPDR. (**B**) Figure depicts increase in average proportions of decisions around the individual decision when viewing concurring cues than when viewing neutral cues. The left part of the figure considers cases when the individual decision was face while the right part considers cases when it was car. The bold dots depict the average across the individuals. (**C**) Mean proportion of decisions towards conflicting cues across individuals. Figure shows that crossover happens for all cases of individual confidence and is most prominent when individual decision confidence is low (5,6). Error bars denote ± SD. (**D**) Social bias ranking of subjects indicating their tendency to be influenced by social cue shown. Larger and darker dots indicate subjects having greater social influence. The dotted line parallel to the x-axis depicts the significance level.

Gaussian distribution has been previously used to model behavioural data successfully [37]. Hence the proposed model was compared with the mixture of two component Gaussian distributions. The median of the likelihood ratios across subjects for a given (*k*_1,_ *k*_2)_ in all but 2 cases (out of 30) clearly indicates that the proposed model outperforms the Gaussian mixture model in terms of explaining the data (refer to Table S3).

#### Effect of Social Cues on Individual Choice

Effect of social cues on individual decision was studied using a distance metric between *k*_1_ and the estimated modes of the fitted model (see equation (6)). Using bootstrap resampling technique on mean distance per (*k*_1,_ *k*_2)_ pair, it can be observed that post social cue, there is a significant shift in ratings when decisions from all subjects were pooled together (Table S4). Furthermore, to check whether this is also true for individual decisions, an additional analysis was carried out. If the proposed model predicted a mode in the direction of the social cue, the proportion of decisions in between *k*_1_ and *k*_2_ was calculated by integrating the estimated density within the said interval. It can be seen that a significant proportion of decisions, as assessed by our model, falls in between *k*_1_ and *k*_2_ (refer to Tables S5–S8), clearly suggesting that, in general, subjects tend to get influenced by the social choice, irrespective of whether it conforms to his/her individual bias or not.

#### Effect of Concurring Cues

In order to check whether the decision confidence increases when the subject was given a social cue which concurs with his/her own judgment, the area under the fitted density given the social cue (‘FF’,’CC’) is compared with that of a neutral cue (‘FC’/’CF’) (see Tables S9 and SlO). These areas are assumed to be indicative of the proportion of decisions of the subjects around the individual decision. As compared to the neutral cue, for most of the subjects the average proportion of decisions in the region [1, 6] is greater when individual choice is face and social cue is also face. Similarly, this proportion in the region [7, 11] is greater when individual and social choice both are car. Thus it can be concluded (refer to Fig. 2B) that decision confidence of most subjects increased when provided with a concurring social cue (FF/CC).

#### Effect of Conflicting Cues

Further analysis was carried out to check whether there is a significant reversal in the decisions when the subject faces a social cue contradictory to his/her individual decision. We say that there is a *cross-over* if there exists a mode in the opposite side of the decision boundary. Cross-over under the influence of concurring cues was found to be insignificant (in terms of area) compared to conflicting cues (see Table S14) and hence ignored. For every *k*_1,_ it is examined whether cross-over exists given a mismatch between social cue and individual choice. Using bootstrapping, it can be shown that the proportion of cross-over is significant among the individuals. This is evident from the approximate achieved significance level (ASL) [63] contained in Table Sll. Fig. 2C distinctly reveals that the mean cross-over proportion increases with decrease in individual confidence, implying that in general subjects tend to be influenced more by contradictory social cues on images where their individual confidence was low. Refer to Tables S12 and S13 for the detailed list of cross-over proportions per subject.

#### Ranking Subjects Based on Social Cues

Individuals differ in the manner in which social information influences their perceptual decision. Using the proposed behavioural model, it is possible to rank the subjects based on the level of influence social cues had on their percept. Fig. 2D shows the ranking of subjects based on a measure, called as social bias score, that captures their tendency to be influenced by the social information. Based on the analysis, 8^1^ subjects were chosen to be most affected by social cues and are referred as *chosen subjects* in the EEG analysis.

### Neural Results

#### ERP Analysis

ERP analysis was performed on average referenced and baseline subtracted EEG signals for each condition. Epochs of a particular channel were marked noisy if their respective absolute differences from the median exceeded 5 times the interquartile range. Such noisy epochs were not considered for further ERP analysis. It is well-known that parieto-occipital electrodes show differential activity when perceiving faces and cars [64]. Several studies have hypothesized the role of the frontal cortex in choice manipulation under the influence of social information ([8, 13, 15, 56]). To explore the effect of social cues on face/car percepts, ERP analysis was carried out with parieto-occipital and fronto-central electrodes separately. To elucidate whether different types of comments induce different neural processing mechanisms, the grand average difference waves were plotted (refer to Fig. 3) for correctly guessed face and car trials. A difference in face and car ERPs is visible across both fronto-central and parieto-occipital electrodes around 200 ms post stimulus onset closely matching the Nl 70 [65] component known to be present in face and non-face ERP s. The difference between concurring and conflicting conditions however seems more prominent around 250-300 ms post stimulus condition in both parieto occipital and fronto central electrodes. Further detailed analysis is carried out using single trial multivariate analysis.

**Figure 3:**
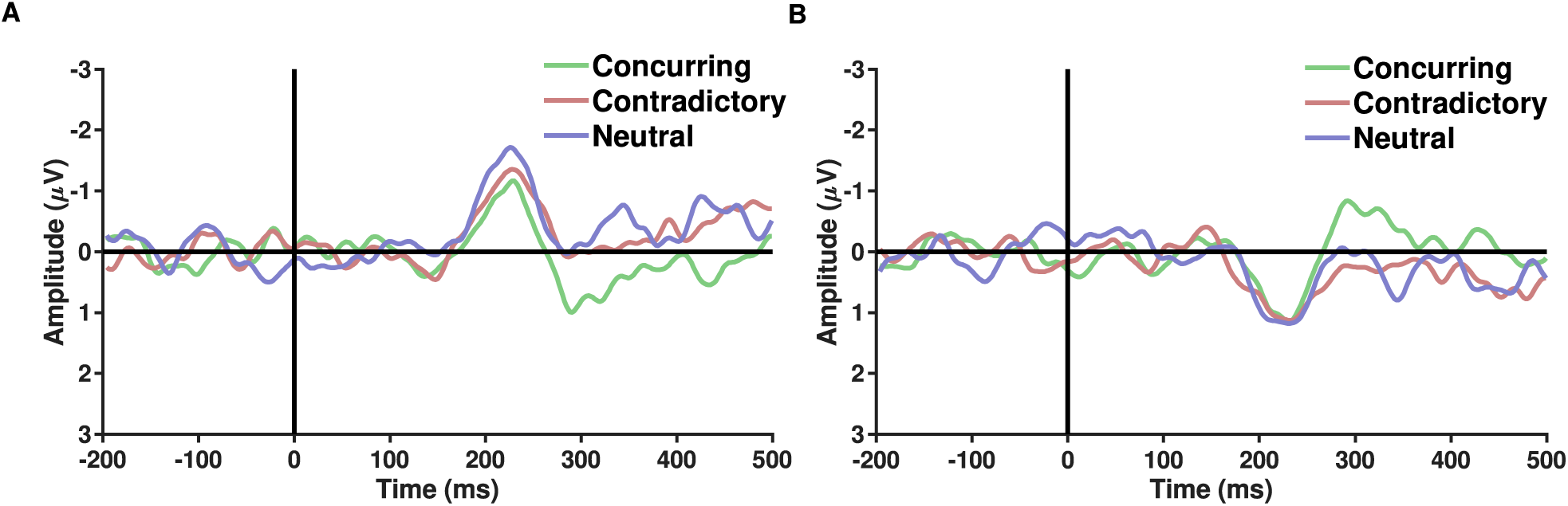
Figures A and B show the grand average of difference ERPs (Face – Car) over parieto occiptial and fronto central electrodes respectively across the three types of conditions - Concurring, Conflicting and Neutral. A sharp peak in the difference waveforms is observed post 200 ms across all conditions. Difference between conflicting and concurring cues seems more prominent around 250-300 ms.

#### Single Trial Multivariate Analysis

Pattern classifiers were used to analyze single trial EEG signals corresponding to the different types of social cues. To quantify the predictive accuracy of the classifier, the posterior probabilities obtained from 10 fold cross validation were used to calculate the area under the ROC curve (AUC). The AUCs were averaged across the subjects. The multivariate analysis was performed using the entire post-stimulus data using all channels and all time points and the AUCs were plotted corresponding to the different conditions(Fig. 4A). The classification accuracy appears to be more while the subject was provided with a cue that concurred with his/her individual guess than while he/she was provided with a conflicting social cue (p= 0.0213,df= 1 4,t= 2.2314). An overall increase in difference was noted between the conditions (p= 0.0038, df= 7,t = 3. 7147, corresponding to the null hypothesis of no difference in the classification rates between the two conditions) when an average over chosen subjects was considered (Fig. 4A). The pattern analysis was executed separately using eeg data for all electrodes across different time windows each having a length of 50 ms. AUCs corresponding to the late sensory period (200-450 ms after stimulus onset) are found to be significantly more than chance (p-value<0.05, false discovery rate (FDR) corrected) for concurring trials. Further analysis shows that the difference between AUCs of concurring and conflicting cues is statistically significant only in the time window 200 – 250 ms (p-value(without multiple correction)= 0.01, t = 2.585, df = 14, FDR corrected p-value=0.054,multiple hypothesis testing performed across time points where the classification rates corresponding to concurringtrials are more than chance). On performing similar time-window wise analysis on the chosen subjects it is seen that the difference stands out to be statistically significant (p-value(without multiple correction)= 2.40 × 10^−5^, t = 8.8377, df = 7, FDR corrected p-value< <0.05) in the 200-250 ms time window.

**Figure 4:**
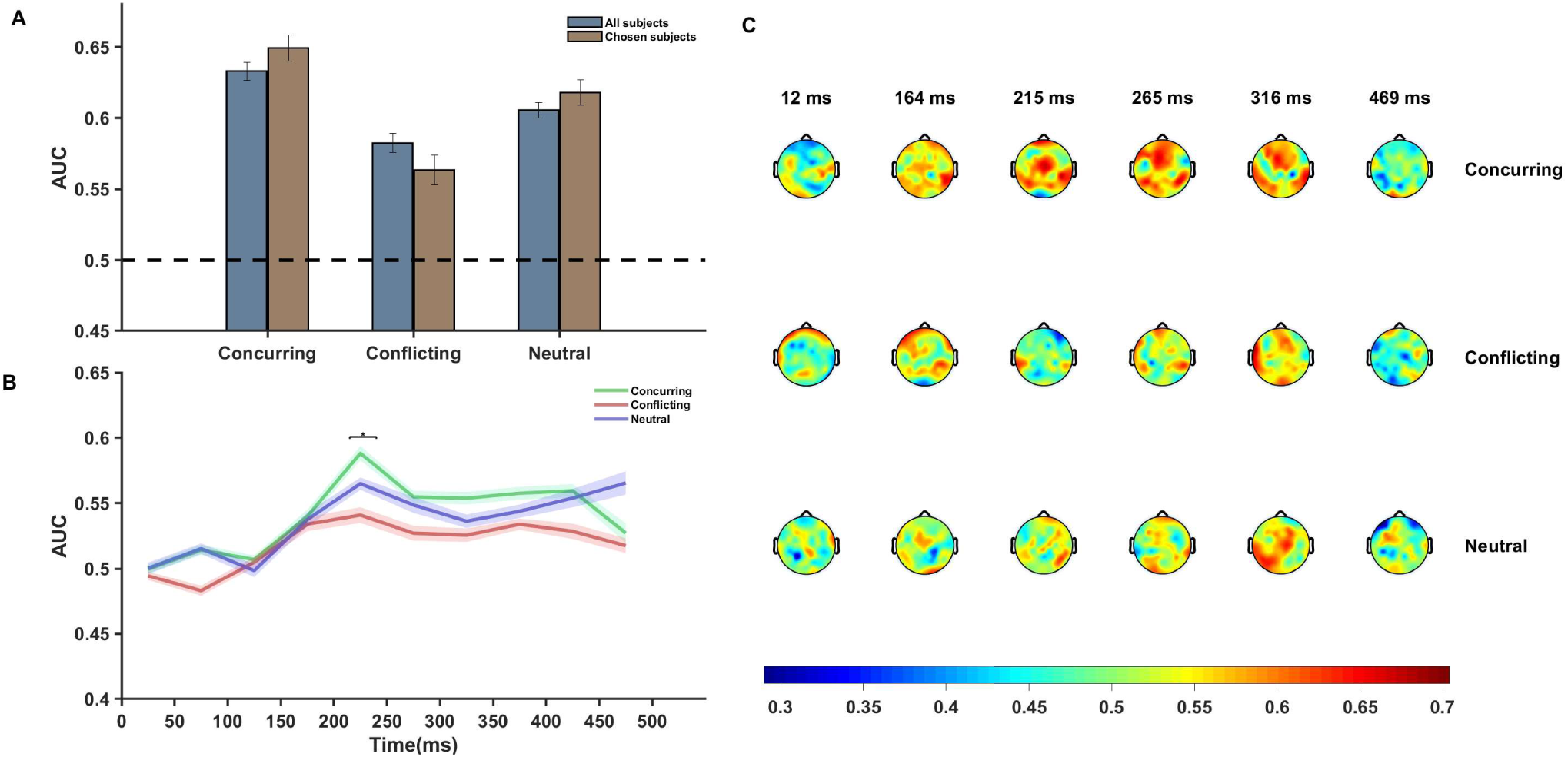
Neural Data Analysis. (A) Figure shows average AUC predicting choice probability using single trial EEG analysis using multivariate pattern analysis. Average AUC increases under the influence of concurring cues and decreases under the influence of conflicting cues as compared to that for neutral cues in all our subjects. The effect is more prominent in case of the chosen subjects. Error Bars indicate± SEM. (B) Plot of average AUC across all subjects at different time points. The increase in AUC is most pronounced in the 200-300 ms post stimulus interval. The difference between the AUCs of concurring and conflicting trials is most significant (p=0.054,FDR corrected) in the 200-250 ms window (marked using *). (C) Topoplot of one subject showing per electrode per time window single trial classification under different cue conditions. Average AUC of the all channels for successive time windows are shown. There appears to be a significant involvement of the frontal and occipital electrodes 200-350 ms post stimuli onset.

Fig. 4B clearly depicts that around 200 - 250 ms after stimulus onset, there is a sharp increase in the AUC value and the peak is more pronounced for concurring social cues. Notably, prominent activity in fronto-central and occipito-temporal electrodes in similar time window was also observed during ERP analysis.

Additional classifier analysis was carried out using data for each electrode separately for each of the time windows(Fig. 4C) and the plot of scalp topography on the basis of the classifier performances (see Fig. 4C) for individual electrodes seems to be consistent with the temporal findings (Fig. 4B). Around 200-300 ms post stimulus onset we observe increased classification accuracy in the parieto-occipital regions and fronto-central regions across all conditions (concurring, conflicting and neutral). In these regions the magnitude of the AUCs are more in case of concurring trials than conflicting and neutral trials (see Fig. 4C). The classifier results demonst rate that social decisions have an effect on individual perceptual decision and it is most prominent around 200-300 ms post stimulus onset.

#### Source Reconstruction Results

Single trial multivariate data analysis and ERP analysis revealed prominent discriminatory activity 200 ms post stimulus onset. Source estimates identified more frontal activity under the influence of conflicting cues than concurring cues (refer to Fig. 5). Frontal sources seem to to be primarily responsible for generating differences in the ERP waveforms of face and car trials across the whole neural timeline for conflicting trials while a prominent frontoparietal interplay was noticed in case of concurring and neutral trials. Particularly, the medial frontal gyrus seems to have contributed significantly in presence of conflicting cues, in line with previous studies which also highlight the role of medial frontal cortex during social conformity and cognitive dissonance [8, 12, 15]. The neural sources of the difference in the current density power between the concurring and conflicting conditions were analyzed using sLORETA with a one-tailed F-ratio test (concurring < conflicting) on paired data separately for 200-250 ms and 250-300ms time windows. Based on the exceedance proportion test [66, 67] results which showed a threshold of 2.38 for a p-value of 0.058 for the 200-250 ms window and a threshold of −2.169 for a p-value of 0.059 for the 250-300 ms window, differences were localized mostly to the frontal areas (refer to tables S16, Sl 7 in SI for the complete list). We found the maximal differences in the medial frontal gyrus (BA 10, MNI coordinates: x = 40, y = 55, z = 0) in both the cases (refer to figures 5E, 5F and tablesS16, Sl 7).

**Figure 5:**
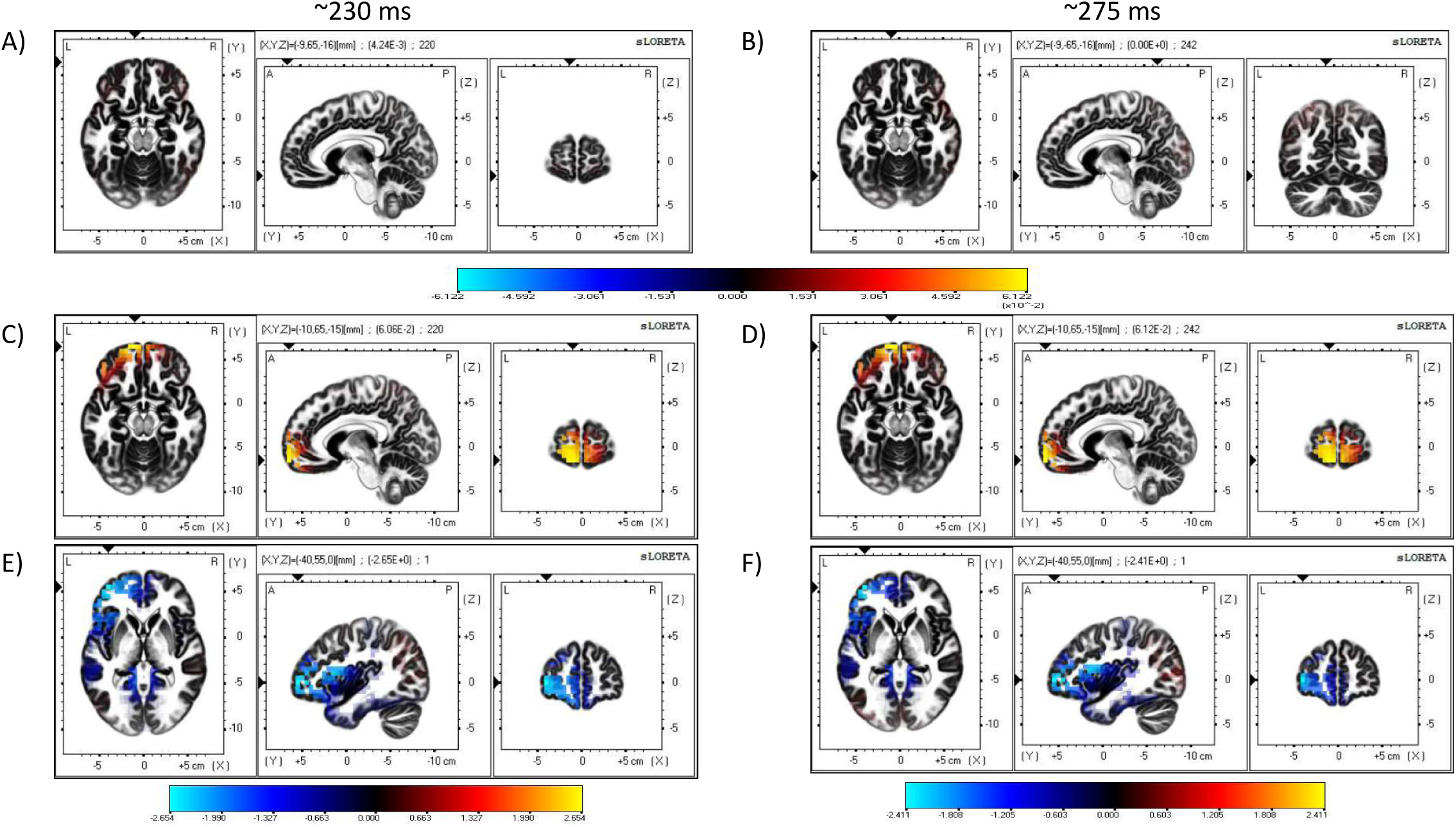
Source Reconstruction. Figures (**A**), (**B**) show sources estimated at 230 ms and 275 ms using sLORETA software for trials with concurring cues. Figures (**C**), (**D**) show sources estimated at 230 ms and 275 ms using sLORETA software for trials with conflicting cues. The color bar depicts the squared magnitude of the current density [*μ A*^2^/(*mm*^4^.H*z*)]. Figures (E) and (F) denote maps of non-parametric statistics comparing concurring and conflicting trials during 200-250ms and 250-300 ms time windows. Non-parametric analysis was performed using one-tailed F-ratio test (concurring < conflicting) on paired data. Color bar represents value of log F-ratio for each voxel.

#### Neural Analysis of Cue Data

We did an additional analysis based on the neural signals when the cue was displayed. We extracted the EEG signals locked to the cue onset. The 500 ms post-cue onset data was used to perform multivariate pattern analysis for exploring the effects of expectation on early sensory processing. If the participants’ response were driven by the cues, then we would expect a higher classification rate for images selected as faces post cue-onset when preceded by an ‘FF’ cue and vice versa for ‘CC’ cues. However, pattern analysis of cue-data revealed no such trends (refer to Fig. 6) and resulted in chance performance for all conditions *(p* > 0.05). A two-way anova was performed to find the statistically significant difference between the four different cue conditions taking into account face and car trials separately along with interactions. The differences are all insignificant (see Table S15) pointing to the fact that there was no significant difference on the classification accuracy across all the cue conditions including the condition where no cue was shown. It is interesting to note that similar chance performance was also observed in pre-stimulus and early post-stimulus (< 200 ms) neural classification. Thus based on the cue analysis, it seems unlikely that the participants’ decision was influenced by cue-based expectation bias in the post cue onset and early visual processing stage following the stimulus display.

**Figure 6:**
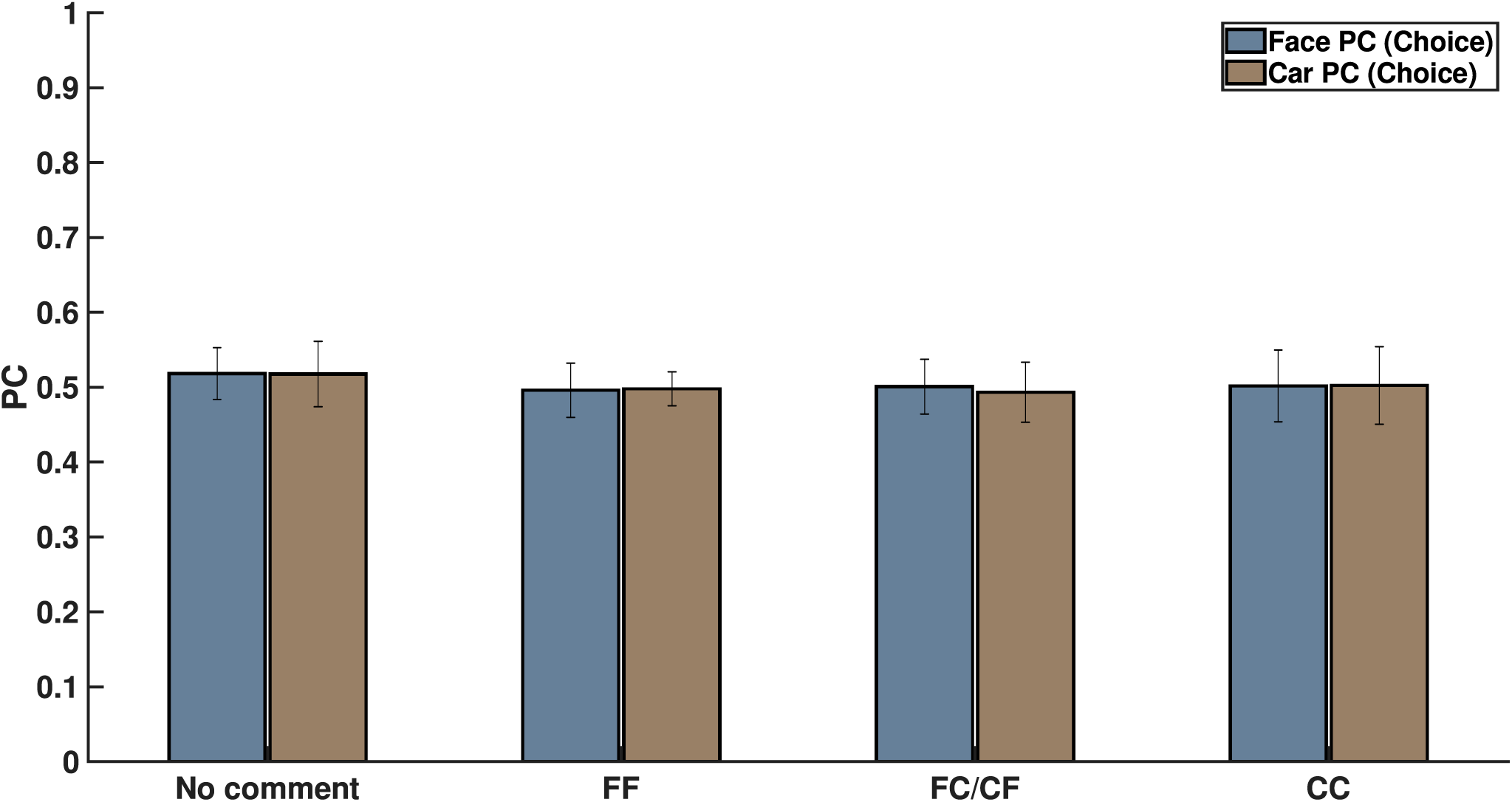
Figure shows percentage of correctly classified face and car decisions for the 4 kinds of comments shown on screen on the basis of their neural signals after cue exposure. This clearly shows that subject choice did not arise from cue-related expectation bias.

## Discussion

How social decision affects individual decision making has been explored in social psychology since 1940’s starting with research on social conformity by Solomon Asch [5, 6, 16] and with the advent of social media, there has been a renewed interest in social cues influencing our decision [1–4]. In the current study, how people respond to social cues when performing a perceptual decision making task was explored systematically. The neural mechanism of the decision making process was studied while the subjects used the social cues in form of two other well performing subjects’ decision, to perceive noisy images of faces and cars. Although the social cues shown to the subject were non-informative with equal number of FF, neutral and CC cues per stimuli displayed in random order, they were found to be successful in manipulating the percept. Most of the studies on social influence require participants to make a decision with and without social cues sequentially but we demonstrate that irrespective of the order in which the stimulus/cue was presented, social cues always have similar effect on our individual decision making. We conclude that the perceptual decision of the subject under the influence of the social cue depends on two factors - his/her individual perception of the image as is reflected in his/her confidence ratings on the same images without any social cue and the social information presented to him/her. It is observed that the distribution of confidence ratings under the influence of a social cue is bi-modal in nature with one mode corresponding to individual decision while other due to social cue (Fig. 1B), with a significant proportion in the direction of the social cue. So we can safely infer that although there was a general tendency to adhere to one’s individual decision, but subjects’ decision confidence could be altered with social influence. This shift in decision confidence varied between the subjects as reported in previous studies [1]. Using the proposed computational model, the heterogeneity of the influence of social cues on the subjects’ decision was quantified successfully. The subjects were ranked based on the influence the social cues elicited and the findings used in subsequent neural analysis produced encouraging results.

Although social influence on perceptual decisions remains a highly researched topic, but the neural mediators of manipulation of perceptual decisions with social influence remains largely unexplored [8, 13, 15, 56]. We identified a sharp peak in the mean AUC value 200-300 ms post stimulus onset which is most prominent in concurring cues. This seems to imply that the classifier could identify the class-specific discriminatory activity and predict the participants decision more accurately when the cue received matched with his/her individual perception, in line with our claim that the subjects were more sure about their decisions when the stimulus was preceded by a concurring cue. The effect is more well-defined in case of car trials, probably arising out of heavier mental load for car images than faces. Humans are adept at face perception [68] and the stimuli displayed had uniform noise for both face and ear thereby making the car detection task comparatively difficult. Figure 2B shows this effect for concurring cues where the increase in decision confidence was more prominent for CC cues than for FF cues.Similar trend is noticed for conflicting cues (Figure 2C) where significant reversal of decision in the direction of the social cues was noticed and proportion of crossover is more for trials originally detected as cars. Almost all the existing neuroimaging studies using social cues suggest the role of posterior medial frontal cortex (pMFC) and to some extent ventral striatum [8, 12, 15] in social conformity but the neural mechanism remains poorly understood. Current research shows that activation in pMFC is modulated by the difference between individual choice and groups’ preference. Role of pMFC in social conformity is further strengthened by a TMS study [13] where participants showed reduced social conformity when pMFC was disrupted. One plausible interpretation of involvement of pFMC could be that conforming to social opinions trigger similar circuitry with that of reinforcement learning [12]. Neural activity in pMFC might mirror activity similar to a prediction error signal which can then be subsequently used to modify or strengthen the perceptual decision. In the current study, source analysis of ERP signals using conflicting cues also shows activity in the medial frontal cortex (MFC) starting around 200 ms post stimulus onset. Neural signals following conflicting cues displayed comparatively greater frontal activity than concurring cues (Figure 5) possibly suggesting greater top down processing of information when cues mismatch perceptual choice. It is particularly interesting to note that MFC is active in the time interval immediately following the well established Nl 70 component which is known to account for difference between face and car [69]. Possibly the mismatch between the top down expectation produced by the social cue and the bottom up sensory information post stimlus onset triggered activity in the MFC which has been reported to play a role in social conformity [12, 18].Medial frontal cortex perhaps generates a signal that encodes the difference between individual percept based on the stimulus and the group decision given by the social cues. Absence of frontal activity in concurrent cues in the same time interval further supports our claim. The strength of MFC activity has been shown to regulate the level of the subsequent adjustment of individual choice [8]. Hence the MFC activation was more pronounced for chosen subjects. Our results seem to suggest that irrespective of stimulus order, neural circuitry similar to existing social conformity studies was shown to be active in making perceptual decision under the influence of social cues.

There has been extensive research on face and object perception in the last few decades revealing significant involvement of various occipito-parietal regions in the early stages of visual processing (< 200 ms) [64]. Additionally, there has been a significant body of work citing that stimulus expectation leads to changes in early sensory processing [24–34]. It has been demonstrated in numerous studies that expectation about stimulus in the form of predicting cues leads to a stimulus bias. Top down expectation effects can be seen in the form of improvement in stimuli representation [29], generation of stimulus template in striate and extrastriate regions [34, 70], and even reduction in amplitude in neural signals leading to “expectation suppression” effect [71]. On the whole, top down expectations in form of predictive cues has been shown to bias neural activity in the pre-stimulus and early sensory processing stage thereby orienting the bottom up sensory information towards one perceptual decision. On the other hand, recent studies have questioned the role of neural expectation in sensory cortex [35, 36]. In our study however, probing into the neural time series unveiled no significant differences in perception under the influence of different social cues during early stages. We systematically analyzed the effect of social decision and found no significant effect of the social cues before stimulus onset, post cue onset and immediately following stimulus onset. We extracted the neural data locked to cue presentation and used multivariate pattern classifier on the cue data alone to show that the cue data were not indicative of any early top down expectation based effect on the stimuli (see Fig. 6). Our results seem to suggest that unlike studies involving predictive cues [72], expectation by virtue of social influence does not affect early sensory processing. It is worthwhile to note here that our cues were essentially social decisions of others instead of cues predictive about the stimulus itself [72, 73] and could possibly explain the lack of top-down expectation signals seen in early sensory cortex in previous studies [72]. Our results seem to suggest that role of downstream processing in using the social information from the cue provided, similar to the concept of Bayesian Decision Theory [74] and Signal Detection Theory [75, 76].

Overall we conclude that perceptual decision and confidence is influenced by social cues and it is possible to compute the extent of influence using statistical modeling. Neural data analysis alludes to a role of a medial frontal cortex affecting perceptual decision under social influence. We found no expectation-related bias in early sensory processing using social information cues. Future studies can possibly focus on experiments using actual social groups to validate the neural results found in the current research.

## Acknowledgment

This work is funded by CSRI-DST, and DST-INSPIRE, Government of India. The authors are thankful to the two anonymous reviewers for many suggestions which improved the article significantly.

## Footnotes

1 Out of the 17 subjects, 2 had only high confidence trials and hence not considered. Out of the 15 remaining, 8 were found to be significantly more affected by the social cues than the rest.

## References

[1] Bertrand Jayles, Hye-rin Kim, Ramón Escobedo, Stéphane Cezera, Adrien Blanchet, Tatsuya Kameda, Clément Sire, and Guy Theraulaz. How social information can improve estimation accuracy in human groups. Proceedings of the National Academy of Sciences, pages 12620–12625, 2017.

[2] Bernard J Jansen, Mimi Zhang, Kate Sobel, and Abdur Chowdury. Twitter power: Tweets as electronic word of mouth. Journal of the American society for information science and technology, 60(11):2169–2188, 2009.

[3] B. Gonçalves and N. Perra. Social Phenomena: From Data Analysis to Models. Computational Social Sciences. Springer International Publishing, 2015. ISBN 9783319140117.

[4] Robert M Bond, Christopher J Fariss, Jason J Jones, Adam DI Kramer, Cameron Marlow, Jaime E Settle, and James H Fowler. A 61-million-person experiment in social influence and political mobilization. Nature, 489(7415):295–298, 2012.

[5] Solomon E Asch. Opinions and social pressure. Scientific American, 193(5):31–35, 1955.

[6] Solomon E Asch and H Guetzkow. Effects of group pressure upon the modification and distortion of judgments. Groups, leadership, and men, pages 222–236, 1951.

[7] GS Berns, J Chappelow, CF Zink, G Fagnoni, ME Martin-Skurski, and J Richards. Neurobiological correlates of social conformity and independence during mental rotation. Neuropsychopharmacology, 29:S77–S77, 2004.

[8] Gregory S Berns, C Monica Capra, Sara Moore, and Charles Noussair. Neural mechanisms of the influence of popularity on adolescent ratings of music. Neurolmage, 2010.

[9] Timothy EJ Behrens, Laurence T Hunt, Mark W Woolrich, and Matthew FS Rush-worth. Associative learning of social value. Nature, 456(7219):245, 2008.

[10] Guido Biele, Jörg Rieskamp, Lea K Krugel, and Hauke R Heekeren. The neural basis of following advice. PLoS biology, 9(6): e1001089, 2011.

[11] Vasily Klucharev, Ale Smidts, and Guillén Fernández. Brain mechanisms of persuasion: how expert powermodulates memory and attitudes. Social cognitive and affective neuroscience, 3(4):353–366, 2008.

[12] Vasily Klucharev, Kaisa Hytönen, Mark Rijpkema, Ale Smidts, and Guillén Fernández. Reinforcement learning signal predicts social conformity. Neuron, 61(1):140–151, 2009.

[13] Vasily Klucharev, Moniek AM Munneke, Ale Smidts, and Guillen Fernandez. Down regulation of the posterior medial frontal cortex prevents social conformity. Journal of Neuroscience, 31(33):11934–11940, 2011.

[14] Daniel K Campbell-Meiklejohn, Dominik R Bach, Andreas Roepstorff, Raymond J Dolan, and Chris D Frith. How the opinion of others affects our valuation of objects. Current Biology, 20(13): 1165-1170, 2010.

[15] Keise Izuma and Ralph Adolphs. Social manipulation of preference in the human brain. Neuron, 78(3):563–573, 2013.

[16] Henri Tajfel. Social psychology of intergroup relations. Annual review of psychology, 33(1):1–39, 1982.

[17] Robert B. Cialdini and Noah J. Goldstein. Social influence: Compliance and conformity. Annual Review of Psychology, 55(1):591–621, 2004. doi:10.1146/annurev.psych.55.090902.142015. PMID: 14744228.

[18] Keise lzuma. The neural basis of social influence and attitude change. Current opinion in neurobiology, 23(3):456–462, 2013.

[19] lain D Couzin. Collective cognition in animal groups. Trends in cognitive sciences, 13 (1):36–43, 2009.

[20] Larissa Conradt and Christian List. Group decisions in humans and animals: a survey. Philosophical Transactions of the Royal Society of London B: Biological sciences, 364 (1518):719–742, 2009.

[21] Andrew M Simons. Many wrongs: the advantage of group navigation. Trends in ecology & evolution, 19(9):453–455, 2004.

[22] Francis Galton. Vox populi. Nature, 75(7):450–451, 1907.

[23] J. Surowiecki. The Wisdom of Crowds. Knopf Doubleday Publishing Group, 2005. ISBN 9780307275059.

[24] Peter Kok, Gijs Joost Brouwer, Marcel AJ van Gerven, and Floris P de Lange. Prior expectations bias sensory representations in visual cortex. Journal of Neuroscience, 33 (41) :16275–16284, 2013.

[25] Maxine T Sherman, Ryota Kanai, Anil K Seth, and Rufin VanRullen. Rhythmic influence of top-down perceptual priors in the phase of prestimulus occipital alpha oscillations. Journal of cognitive neuroscience, 28(9):1318–1330, 2016.

[26] Ana Todorovic, Jan-Mathijs Schoffelen, Freek van Ede, Eric Maris, and Floris P de Lange. Temporal expectation and attention jointly modulate auditory oscillatory activity in the beta band. PLoS One, 10(3):e0120288, 2015.

[27] Elexa St John-Saaltink, Christian Utzerath, Peter Kok, Hakwan C Lau, and Floris P De Lange. Expectation suppression in early visual cortex depends on task set. PLoS One, 10(6):e0131172, 2015.

[28] Peter Kok, Dobromir Rahnev, Janneke FM Jehee, Hakwan C Lau, and Floris P De Lange. Attention reverses the effect of prediction in silencing sensory signals. Cere bral cortex, 22(9):2197–2206, 2012.

[29] Peter Kok, Janneke FM Jehee, and Floris P De Lange. Less is more: expectation sharpens representations in the primary visual cortex. Neuron, 75(2):265–270, 2012.

[30] .Jiefeng Jiang, Christopher Summerfield, and Tobias Egner. Attention sharpens the distinction between expected and unexpected percepts in the visual brain. Journal of Neuroscience, 33(47):18438–18447, 2013.

[31] Katrina Carlsson, Predrag Petrovic, Stefan Skare, Karl Magnus Petersson, and Martin Ingvar. Tickling expectations: neural processing in anticipation of a sensory stimulus. Journal of cognitive neuroscience, 12(4):691–703, 2000.

[32] Peter Kok, Lieke Lf Van Lieshout, and Floris P De Lange. Local expectation violations result in global activity gain in primary visual cortex. Scientific reports, 6:37706, 2016.

[33] Peter Kok, Pim Mostert, and Floris P De Lange. Prior expectations induce prestimulus sensory templates. Proceedings of the National Academy of Sciences, page 201705652, 2017.

[34] Peter Kok, Michel F Failing, and Floris P de Lange. Prior expectations evoke stimulus templates in the primary visual cortex. Journal of Cognitive Neuroscience, 26(7):1546-1554, 2014.

[35] Nuttida Rungratsameetaweemana, Sirawaj Itthipuripat, Annalisa Salazar, and John T Serences. Expectations do not alter early sensory processing during perceptual decision making. Journal of Neuroscience, pages 3638–17, 2018.

[36] Ji Won Bang and Dobromir Rahnev. Stimulus expectation alters decision criterion but not sensory signal in perceptual decision making. Scientific reports, 7(1):17072, 2017.

[37] Seongmin A Park, Sidney Goïame, David A O’Connor, and Jean-Claude Dreher. Integration of individual and social information for decision-making in groups of different sizes. PLoS Biology, 15(6):1–28, 2017.

[38] Nikolaus F Troje and Heinrich H Biilthoff. Face recognition under varying poses: The role of texture and shape. Vision research, 36(12):1761-1772, 1996.

[39] Charalampos Chanialidis. Bayesian mixture models for count data. PhD thesis, University of Glasgow, 2015.

[40] G. Casella and R.L. Berger. Statistical Inference. Duxbury advanced series in statistics and decision sciences. Thomson Learning, 2002. ISBN 9780534243128.

[41] K.E. Atkinson. An introduction to numerical analysis. Wiley, 1978. ISBN 9780471029854.

[42] J. D. Gibbons and S. Chakraborti. Nonparametric Statistical In ference. Taylor & Francis, 2011. ISBN 9781420077612 - CAT C7619.

[43] P.H. Westfall, P.H.W. S. Stanley Young, and S.S. Young. Resampling-B ased Multiple Testing: Examples and Methods for P-Value Adjustment. A Wiley-Interscience publication. Wiley, 1993. ISBN 9780471557616.

[44] Rob J Hyndman. Computing and graphing highest density regions. The American Statistician, 50(2):120–126, 1996.

[45] Koel Das, Barry Giesbrecht, and Miguel P Eckste in. Predicting variations of perceptual performance across individuals from neural activity using pattern classifiers. Neuroim age, 51(4):1425 –1437, 2010.

[46] Yukiyasu Kamitani and Frank Tong. Decoding the visual and subjective contents of the human brain. Nature neuroscience, 8(5):679, 2005.

[47] Marios G Philiastides, Roger Ratcliff, and Paul Sajda. Neural representation of task difficulty and decision making during perceptual categorization: a timing diagram. Journal of Neuroscience, 26(35):8965–8975, 2006.

[48] John-Dylan Haynes and Geraint Rees. Predicting the orientation of invisible stimuli from activity in human primary visual cortex. Nature neuroscience, 8(5):686, 2005.

[49] Koel Das and Zoran Nenadic. An efficient discriminant-based solution for small sample size problem. Pattern Recognition, 42:857–866, 2009.

[50] Koel Das, Daniel S Rizzuto, and Zoran Nenadic. Mental state estimation for brain computer interfaces. IEEE Transactions on Biomedical Engineering, 56(8):2114–2122, 2009.

[51] An H Do, Po T Wang, Christine E King, Ahmad Abiri, and Zoran Nenadic. Brain computer interface controlled functional electrical stimulation system for ankle movement. Journal of neuroengineering and rehabilitation, 8(1):49, 2011.

[52] An H Do, Po T Wang, Christine E King, Sophia N Chun, and Zoran Nenadic. Brain computer interface controlled robotic gait orthosis. Journal of neuroengineering and rehabilitation, 10(1):111, 2013.

[53] Christine E King, Po T Wang, Luis A Chui, An H Do, and Zoran Nenadic. Operation of a brain-computer interface walking simulator for individuals with spinal cord injury. Journal of neuroengineering and rehabilitation, 10(1):77, 2013.

[54] Po T Wang, Christine E King, Luis A Chui, An H Do, and Zoran Nenadic. Self-paced brain-computer interface control of ambulation in a virtual reality environment. Journal of neural engineering, 9(5):056016, 2012.

[55] Richard O Duda, Peter E Hart, and David G Stork. Pattern classification. John Wiley & Sons, 2012.

[56] Simon J Mason and Nicholas E Graham. Areas beneath the relative operating characteristics (roe) and relative operating levels (rol) curves: Statistical significance and interpretation. Quarterly Journal of the Royal Meteorological Society, 128(584):2145–2166, 2002.

[57] Roberto Domingo Pascual-Marqui et al. Standardized low-resolution brain electromagnetic tomography (sloreta): technical details. Methods Find Exp Clin Pharmacol, 24 (Suppl D):5–12, 2002.

[58] Manfred Fuchs, Jörn Kastner, Michael Wagner, Susan Hawes, and John S Ebersole. A standardized boundary element method volume conductor model. Clinical Neurophysiology, 113(5):702 –712, 2002.

[59] John Mazziotta, Arthur Toga, Alan Evans, Peter Fox, Jack Lancaster, Karl Zilles, Roger Woods, Tomas Paus, Gregory Simpson, Bruce Pike, et al. A probabilistic atlas and reference system for the human brain: International consortium for brain mapping (icbm). Philosophical Transactions of the Royal Society of London B: Biological sciences, 356(1412):1293–1322, 2001.

[60] J Talairach and P Tournoux. Co-planar stereotaxic atlas of the human brain: 3-d proportional system: An approach to cerebral imaging (thieme classics). Thieme, 1988.

[61] Jack L Lancaster, Marty G Woldorff, Lawrence M Parsons, Mario Liotti, Catarina S Freitas, Lacy Rainey, Peter V Kochunov, Dan Nickerson, Shawn A Mikiten, and Peter T Fox. Automated talairach atlas labels for functional brain mapping. Human brain mapping, 10(3):120–131, 2000.

[62] Thomas E Nichols and Andrew P Holmes. Nonparametric permutation tests for functional neuroimaging: a primer with examples. Human brain mapping, 15(1):1–25, 2002.

[63] B. Efron and R.J. Tibshirani. An Introduction to the Bootstrap. Chapman & Hall/CRC Monographs on Statistics & Applied Probability. Taylor & Francis, 1994. ISBN 9780412042317.

[64] Bruno Rossion, Carrie A Joyce, Garrison W Cottrell, and Michael J Tarr. Early lateralization and orientation tuning for face, word, and object processing in the visual cortex. Neuroimage, 20(3):1609–1624, 2003.

[65] Shlomo Bentin, Truett Allison, Aina Puce, Erik Perez, and Gregory McCarthy. Electro physiological studies of face perception in humans. Journal of cognitive neuroscience, 8(6):551–565, 1996.

[66] KJ Friston, CD Frith, PF Liddle, Raymond J Dolan, AA Lammertsma, and RSJ Frack owiak. The relationship between global and local changes in pet scans. Journal of Cerebral Blood Flow & Metabolism, 10(4):458–466, 1990.

[67] Karl J Friston, CD Frith, PF Liddle, and RSJ Frackowiak. Comparing functional (pet) images: the assessment of significant change. Journal of Cerebral Blood Flow & Metabolism, 11(4):690 –699, 1991.

[68] David A Leopold and Gillian Rhodes. A comparative view of face perception. Journal of Comparative Psychology, 124(3):233–251, 2010.

[69] Sharon Daniel and Shlomo Bentin. Age-related changes in processing faces from detection to identification: Erp evidence. Neurobiology of Aging, 33(1):206–el, 2012.

[70] Amrita M Puri, Ewa Wojciulik, and Charan Ranganath. Category expectation modulates baseline and stimulus-evoked activity in human inferotemporal cortex. Brain research, 1301:89–99, 2009.

[71] Ana Todorovic and Floris P de Lange. Repetition suppression and expectation suppression are dissociable in time in early auditory evoked fields. Journal of Neuroscience, 32 (39):13389–13395, 2012.

[72] Christopher Summerfield and Floris P De Lange. Expectation in perceptual decision making: neural and computational mechanisms. Nature Reviews Neuroscience, 15(11): 745–756, 2014.

[73] Christopher Summerfield and Etienne Koechlin. A neural representation of prior infor mation during perceptual inference. Neuron, 59(2):336–347, 2008.

[74] Laurence T Maloney and Pascal Mamassian. Bayesian decision theory as a model of human visual perception: Testing bayesian transfer. Visual neuroscience, 26(1):147–155, 2009.

[75] D.M. Green and J.A. Swets. Signal Detection Theory and Psychophysics. Peninsula Pub., 1988. ISBN 9780932146236.

[76] Neil A Macmillan and C Douglas Creelman. Detection theory: A user’s guide. Psychology press, 2004.

