## Supporting Information for "Neural Mediators of Altered Perceptual Choice and Confidence Using Social Information"

### Supplementary Information

#### Calculation of Cumulative Distribution Function

The cdf $G(\cdot)$ of the shifted gamma distribution (equation (1)) is given by $G(y) = 0,$ if $y \leqslant 1$, otherwise,

$$\begin{aligned} G(y) &= \int_{-\infty}^{y} \frac{\beta^{\alpha}}{\Gamma(\alpha)}(x-1)^{(\alpha-1)}e^{-\beta(x-1)}I_{[1,\infty)}(x)\,dx \\ &= \int_{1}^{y} \frac{\beta^{\alpha}}{\Gamma(\alpha)}(x-1)^{(\alpha-1)}e^{-\beta(x-1)}\,dx \\ &= \int_{0}^{y-1} \frac{\beta^{\alpha}}{\Gamma(\alpha)}(z)^{(\alpha-1)}e^{-\beta(z)}\,dz \\ &= \Gamma\left(y-1,\alpha,\frac{1}{\beta}\right). \end{aligned}$$

The cdf $Ng(\cdot)$ of the negative gamma distribution (equation (2)) for $y \leqslant L$ is given by

$$\begin{aligned} Ng(y) &= \int_{-\infty}^{y} \frac{\beta^{\alpha}}{\Gamma(\alpha)}(L-x)^{(\alpha-1)}e^{-\beta(L-x)}I_{(-\infty,L]}(x)\,dx \\ &= \int_{L-y}^{\infty} \frac{\beta^{\alpha}}{\Gamma(\alpha)}z^{(\alpha-1)}e^{-\beta(z)}\,dz \\ &= 1-\Gamma\left(L-y,\alpha,\frac{1}{\beta}\right), \end{aligned}$$

and $Ng(y) = 1$, otherwise. Here

$$I_A(x) = \begin{cases} 1 & \text{if } x \in A, \\ 0 & \text{otherwise,} \end{cases}$$

and $\Gamma(x, \alpha, \frac{1}{\beta})$ is the cdf of the standard gamma distribution with shape parameter $\alpha$ and scale parameter $1/\beta$.

#### Calculation of Mode

It can be easily shown that the mode of the shifted gamma distribution is given by

$$M_g = 1 + \frac{\alpha - 1}{\beta}.$$

For finding the mode of the negative gamma distribution we start by taking the logarithm of its density (equation (2)),

$$\ln(ng(x)) = \alpha \ln(\beta) - \ln(\Gamma(\alpha)) + (\alpha - 1)\ln(L - y) - \beta(L - y).$$

Differentiating with respect to $x$ and equating to zero, we get the mode to be

$$L - \frac{\alpha - 1}{\beta}.$$

#### Calculation of MLE of the parameters of the proposed mixture models

The calculation of MLE of the parameters based on EM algorithm for the mixture of a shifted gamma and a negative gamma model (equation [5]) is demonstrated here. Suppose $X_1, \ldots, X_n$ be i.i.d. observations from

$$f(x) = p\ g_{\alpha_1,\beta_1}(x) + (1-p)\ ng_{\alpha_2,\beta_2}(x).$$

We define an auxiliary variable $Y_i$ such that

$$Y_i = \begin{cases} 1 & \text{if } X_i \sim g_{\alpha_1,\beta_1}(x), \\ 0 & \text{if } X_i \sim ng_{\alpha_2,\beta_2}(x). \end{cases}$$

So the complete likelihood and complete log-likelihood are given by

$$L = \prod_{i=1}^{n} [p\, g_{\alpha_1,\beta_1}(x_i)]^{y_i} [(1-p)\, ng_{\alpha_2,\beta_2}(x_i)]^{(1-y_i)} \text{ and}$$

$$l = \sum_{i=1}^{n} [y_i \ln(p\, g_{\alpha_1,\beta_1}(x_i)) + (1-y_i)\ln((1-p)\, ng_{\alpha_2,\beta_2}(x_i))]\,,$$

respectively. The calculations are done using E-step and M-step.

#### E-step

$y_i$s' are replaced with their conditional expected values

$$\begin{aligned} \widehat{Y_i} &:= E(Y_i|X_i = x_i) \\ &= \frac{p\, g_{\alpha_1,\beta_1}(x)}{p\, g_{\alpha_1,\beta_1}(x_i) + (1-p)\, ng_{\alpha_2,\beta_2}(x_i)}. \end{aligned}$$

#### M-step

The MLE of $p$ is obtained by differentiating $l$ with respect to $p$ and replacing the unobserved $y_i$ by $\widehat{y_i}$ as

$$\hat{p} = \frac{\sum_{i=1}^{n} \widehat{y_i}}{n}.$$

Differentiating $l$ with respect to $\alpha_1$ and $\beta_1$, respectively, and replacing the unobserved $y_i$ by $\widehat{y_i}$, we obtain

$$\frac{\Gamma'(\alpha_1)}{\Gamma(\alpha_1)} - \ln(\alpha_1) = \ln C_1 + \frac{\sum_{i=1}^{n} \widehat{y_i}\ln(x_i - 1)}{\sum_{i=1}^{n} \widehat{y_i}}, \tag{7}$$

and $\beta_1 = \alpha_1 C_1$, where $\Gamma(\cdot)$ is the gamma function and $\Gamma'(\cdot)$ is it's derivative and

$$C_1 = \frac{\sum_{i=1}^{n} \widehat{y_i}}{\sum_{i=1}^{n} \widehat{y_i}(x_i - 1)}.$$

Notice that the closed form solution for the MLE of shape parameter $\alpha_1$ is not tractable. So we use Newton-Raphson technique for getting the numerical solution of the equation (7). Once the MLE of $\alpha_1$ is obtained as $\widehat{\alpha}_1$ using numerical technique, MLE of $\beta_1$, $\widehat{\beta}_1$, is obtained by replacing $\alpha_1$ with $\widehat{\alpha}_1$ in the equation $\beta_1 = \alpha_1 C_1$. Similarly we differentiate $l$ with respect to $\alpha_2$ and $\beta_2$, respectively, and replace the the unobserved $y_i$'s by $\widehat{y_i}$'s to get

$$\ln\alpha_2 - \frac{\Gamma'(\alpha_2)}{\Gamma(\alpha_2)} = \ln\frac{1}{C_2} - \frac{\sum_{i=1}^{n}(1-\widehat{y_i})\ln(L - x_i)}{\sum_{i=1}^{n}(1-\widehat{y_i})} \tag{8}$$

and $\beta_2 = \alpha_2 C_2$, where

$$C_2 = \frac{\sum_{i=1}^{n}(1-\widehat{y_i})}{\sum_{i=1}^{n}(1-\widehat{y_i})(L - x_i)}.$$

As above Newton-Raphson technique is employed to get MLE of $\alpha_2$ as $\widehat{\alpha}_2$ and once it is obtained, the MLE of $\beta_2$ is found by replacing $\alpha_2$ with $\widehat{\alpha}_2$ in the equation $\beta_2 = \alpha_2 C_2$. We start with an initial guess for the parameters $(\alpha_1, \beta_1, \alpha_2, \beta_2)$ and $p$ and then follow the E-step and M-step, iteratively, until convergence.

For the other models given by equations (3) and (4), similar steps as described above have been followed with little modifications. Hence detailed calculations for the other models are omitted here.

### One Sample Hypothesis Testing using Bootstrap

Suppose we want to test the null hypothesis ($H_0$) about a parameter $\theta$ of the distribution $F$ based on a random sample $x_1, \ldots, x_n$. Further assume that the statistical test is done based on a test statistic $T$, measuring the discrepancy between the data and the null hypothesis, such that large values of $T$ indicating evidence against $H_0$. Let the observed value of statistic be given by $t$. Then the achieved significance level is defined as

$$\text{ASL} = P(T \geqslant t | H_0).$$

We estimate the ASL using bootstrap resampling technique. Small value of ASL show the evidence against the null hypothesis.

Table S1: Cases per subject where the proposed model was rejected using KS test (using Holm-Bonferroni correction) with FWER 0.05

| Subject | p-value | $k_1$ | $k_2$ |
|---|---|---|---|
| 1 | 0.0011 | 10 | 10 |
| 2 | — | — | — |
| 3 | 0.0000 | 10 | 10 |
| | 0.0000 | 1 | 1 |
| | 0.0384 | 10 | 5 |
| 4 | 0.0000 | 10 | 10 |
| | 0.0000 | 1 | 1 |
| | 0.0143 | 1 | 10 |
| 5 | — | — | — |
| 6 | — | — | — |
| 7 | 0.0011 | 1 | 1 |
| 8 | — | — | — |
| 9 | — | — | — |
| 10 | 0.0000 | 10 | 10 |
| | 0.0004 | 5 | 5 |
| 11 | 0.0000 | 1 | 1 |
| | 0.0000 | 10 | 10 |
| 12 | — | — | — |
| 13 | — | — | — |
| 14 | — | — | — |
| 15 | — | — | — |
| 16 | 0.0000 | 1 | 1 |
| | 0.0000 | 10 | 10 |
| 17 | — | — | — |

The subjects are marked as '—' when the KS test failed to reject any of its cases. Clearly in more than 96% of the cases our proposed models are accepted.

Table S2: Mean prediction error rate across subjects using 10-fold cross validation

| $k_1$ \ $k_2$ | 1 | 5 | 10 |
|---|---|---|---|
| 1 | 0.009 | 0.0060 | 0.0077 |
| 2 | 0.0119 | 0.0086 | 0.0042 |
| 3 | 0.0055 | 0.0037 | 0 |
| 4 | 0 | 0.0309 | 0.0167 |
| 5 | 0.00625 | 0 | 0.0309 |
| 6 | 0.0077 | 0.0026 | 0.0179 |
| 7 | 0.0104 | 0.0073 | 0.0125 |
| 8 | 0.0370 | 0.030 | 0.0037 |
| 9 | 0.0214 | 0.0119 | 0.0048 |
| 10 | 0.0096 | 0.0026 | 0.0067 |

As evident from the table, the mean prediction error rate never exceeds 5%.

Table S3: Median of likelihood ratio between the proposed model and the Gaussian mixture model across subjects

| $k_1$ \ $k_2$ | 1 | 5 | 10 |
|---|---|---|---|
| 1 | 173728.1573 | 3.3977 | 6.6146 |
| 2 | 2.7082 | 2.8670 | 4.6755 |
| 3 | 2.5848 | 1.6723 | 1.5284 |
| 4 | 1.0451 | 1.9498 | 1.4990 |
| 5 | 1.4201 | **0.9932** | 2.0759 |
| 6 | 1.2763 | 1.3454 | 1.3324 |
| 7 | 1.0303 | **0.9708** | 1.7591 |
| 8 | 1.0961 | 1.4150 | 1.1443 |
| 9 | 2.7505 | 1.8096 | 1.5784 |
| 10 | 28.7567 | 8.0835 | 288.6886 |

In all but 2 (marked in bold) cases our model surpasses the Gaussian mixture model by a clear margin.

Table S4: Approximate achieved significance level (ASL) for testing the shift from individual decision across the subjects

| $k_1$ \ $k_2$ | 1 | 5 | 10 |
|---|---|---|---|
| 1 | **0.091** | 0 | 0 |
| 2 | 0 | 0 | 0 |
| 3 | 0 | 0 | 0 |
| 4 | 0 | 0 | 0.003 |
| 5 | 0 | 0.005 | 0 |
| 6 | 0 | 0.003 | 0 |
| 7 | 0.001 | 0.001 | 0 |
| 8 | 0 | 0 | 0.002 |
| 9 | 0 | 0 | 0.003 |
| 10 | 0 | 0 | 0 |

Lower the ASL more the evidence against the null hypothesis that there is no shift from the individual decision under the influence of a social cue. It is clear that in all but one case (marked in bold) the ASL is less than 0.05.

Table S5: Proportion of decisions between $k_1$ and $k_2$ under the fitted model, for $k_1 \in \{3, 4, 5\}$ and $k_2 = 1$

| sub \ $k_1$ | 3 | 4 | 5 |
|---|---|---|---|
| 1 | 0.6016 | — | — |
| 2 | — | — | — |
| 3 | — | — | — |
| 4 | — | — | 0.7984 |
| 5 | 0.4462 | 0.1880 | 0.3729 |
| 6 | 0.1731 | 0.4863 | 0.5574 |
| 7 | — | 0.6340 | 0.3930 |
| 8 | — | — | 0.4038 |
| 9 | 0.3790 | 0.5146 | 0.6252 |
| 10 | — | — | 0.5781 |
| 11 | — | — | 0.2888 |
| 12 | 0.5393 | 0.4024 | — |
| 13 | 0.7808 | — | 0.9219 |
| 14 | 0.3700 | 0.5984 | 0.6185 |
| 15 | 0.3991 | 0.4726 | 0.3306 |
| 16 | — | — | 0.4002 |
| 17 | 0.6201 | — | — |

The cases marked '—' were not considered due to very few observations for that $(k_1, k_2)$ pair. In all other cases, there was the existence of a mode in the direction of $k_2$.

Table S6: Proportion of decisions between $k_1$ and $k_2$ under the fitted model, for $k_1 \in \{6, 7, 8\}$ and $k_2 = 10$

| sub \ $k_1$ | 6 | 7 | 8 |
|---|---|---|---|
| 1 | — | — | 0.6635 |
| 2 | — | — | — |
| 3 | — | — | — |
| 4 | 0.8245 | — | — |
| 5 | 0.6108 | 0.5679 | 0.3386 |
| 6 | 0.7428 | 0.5922 | 0.3721 |
| 7 | 0.7776 | 0.7090 | — |
| 8 | 0.4759 | — | 0.6070 |
| 9 | 0.6512 | 0.4618 | 0.2916 |
| 10 | 0.6248 | — | — |
| 11 | 0.5959 | — | — |
| 12 | 0.5634 | 0.6094 | 0.6686 |
| 13 | 0.9534 | 0.8441 | 0.8166 |
| 14 | 0.1802 | — | — |
| 15 | 0.7409 | 0.7214 | 0.6222 |
| 16 | — | — | — |
| 17 | 0.7682 | 0.7024 | 0.6154 |

The cases marked '—' were not considered due to very few observations for that $(k_1, k_2)$ pair. In all other cases, there was the existence of a mode in the direction of $k_2$.

Table S7: Proportion of decisions between $k_1$ and $k_2$ under the fitted model, for $k_1 \in \{3, 4, 5\}$ and $k_2 = 10$

| sub \ $k_1$ | 6 | 7 | 8 |
|---|---|---|---|
| 1 | 0.7293 | — | — |
| 2 | — | — | — |
| 3 | — | — | — |
| 4 | — | — | 0.6540 |
| 5 | 0.6978 | 0.7478 | 0.6376 |
| 6 | 0.8773 | 0.6774 | 0.4338 |
| 7 | — | 0.0973 | 0.6147 |
| 8 | — | — | 0.6258 |
| 9 | 0.5831 | 0.7430 | 0.5733 |
| 10 | — | — | 0.7455 |
| 11 | — | — | 0.5767 |
| 12 | 0.5526 | 0.5739 | — |
| 13 | 0.7632 | — | 0.9073 |
| 14 | 0.7670 | 0.5751 | 0.2323 |
| 15 | 0.7586 | 0.6120 | 0.7291 |
| 16 | — | — | 0.4662 |
| 17 | 0.4062 | — | — |

The cases marked '—' were not considered due to very few observations for that $(k_1, k_2)$ pair. In all other cases, there was the existence of a mode in the direction of $k_2$.

Table S8: Proportion of decisions between $k_1$ and $k_2$ under the fitted model, for $k_1 \in \{6, 7, 8\}$ and $k_2 = 1$

| sub \ $k_1$ | 6 | 7 | 8 |
|---|---|---|---|
| 1 | — | — | 0.5862 |
| 2 | — | — | — |
| 3 | — | — | — |
| 4 | 0.4117 | — | — |
| 5 | 0.5217 | 0.4926 | 0.3471 |
| 6 | 0.3266 | 0.5007 | 0.5406 |
| 7 | 0.4153 | 0.4884 | — |
| 8 | 0.4310 | — | 0.2739 |
| 9 | 0.5610 | 0.6997 | 0.6866 |
| 10 | 0.5157 | — | — |
| 11 | 0.3344 | — | — |
| 12 | 0.6305 | 0.6511 | 0.7581 |
| 13 | 0.8309 | 0.6973 | 0.5820 |
| 14 | 0.7860 | — | — |
| 15 | 0.3965 | 0.4116 | 0.4890 |
| 16 | — | — | — |
| 17 | 0.4541 | 0.4347 | 0.4451 |

The cases marked '—' were not considered due to very few observations for that $(k_1, k_2)$ pair. In all other cases, there was the existence of a mode in the direction of $k_2$.

Table S9: Average proportions of decisions within the region [1,6] when the individual choice was face, given $k_2 = 1$ vs $k_2 = 5$

| sub \ $k_2$ | 1 | 5 |
|---|---|---|
| 1 | **0.7511** | **0.6839** |
| 2 | **0.9983** | **0.7982** |
| 3 | **0.9270** | **0.8535** |
| 4 | **0.8141** | **0.6150** |
| 5 | 0.7853 | 0.7983 |
| 6 | **0.8001** | **0.7915** |
| 7 | 0.7163 | 0.7931 |
| 8 | **0.8189** | **0.7981** |
| 9 | **0.7934** | **0.6938** |
| 10 | **0.7791** | **0.6644** |
| 11 | 0.6761 | 0.6955 |
| 12 | **0.8485** | **0.7794** |
| 13 | **0.9784** | **0.8064** |
| 14 | 0.9071 | 0.9145 |
| 15 | 0.7771 | 0.8108 |
| 16 | **0.6699** | **0.6773** |
| 17 | **0.9376** | **0.9216** |

The subjects whose proportion of decisions increased in case of 'FF' cue relative to 'FC/CF' are marked in bold

Table S10: Average proportions of decisions within the region [7,11] when the individual choice was car, given $k_2 = 10$ vs $k_2 = 5$

| sub \ $k_2$ | 10 | 5 |
|---|---|---|
| 1 | **0.7802** | **0.6995** |
| 2 | **0.9909** | **0.7918** |
| 3 | **0.9483** | **0.8235** |
| 4 | **0.8390** | **0.6934** |
| 5 | **0.6971** | **0.6330** |
| 6 | 0.8142 | 0.8359 |
| 7 | **0.8587** | **0.8200** |
| 8 | 0.7937 | 0.8131 |
| 9 | **0.6047** | **0.4368** |
| 10 | **0.7343** | **0.5576** |
| 11 | **0.7627** | **0.7436** |
| 12 | **0.8135** | **0.7581** |
| 13 | **0.9291** | **0.8744** |
| 14 | **0.1802** | **0.1543** |
| 15 | **0.8683** | **0.7740** |
| 16 | 0.8444 | 0.8461 |
| 17 | **0.8828** | **0.7817** |

The subjects whose proportion of decisions increased in case of 'CC' cue relative to 'FC/CF' are marked in bold

Table S11: Approximate achieved significance level (ASL) for testing the cross-over from individual decision towards contradictory social cue

| $k_1$ \ $k_2$ | 1 | 5 | 10 |
|---|---|---|---|
| 1 | | | 0 |
| 2 | | | 0 |
| 3 | | | 0 |
| 4 | | | 0 |
| 5 | | | 0 |
| 6 | 0 | | |
| 7 | 0 | | |
| 8 | **0.1220** | | |
| 9 | 0 | | |
| 10 | 0 | | |

Lower the ASL more the evidence against the null hypothesis that the proportion of subjects having a crossover is not more than 0.5. It is clear that in all but one case (marked in bold) the ASL is less than 0.05.

Table S12: Cross-over area under the influence of conflicting cue 'CC'

| sub \ $(k_1, k_2)$ | (1,10) | (2,10) | (3,10) | (4,10) | (5,10) |
|---|---|---|---|---|---|
| 1 | 0.3251 | 0.6484 | 0.6529 | — | — |
| 2 | 0.1526 | 0.2324 | — | — | — |
| 3 | 0.2711 | 0.4388 | — | — | — |
| 4 | 0.5984 | — | — | — | 0.6361 |
| 5 | nco | 0.1659 | 0.1983 | 0.3208 | 0.4485 |
| 6 | — | 0.1442 | 0.2044 | 0.2887 | 0.3838 |
| 7 | 0.0850 | 0.1460 | — | 0.0911 | 0.5606 |
| 8 | 0.0348 | 0.0800 | — | — | 0.5501 |
| 9 | 0.2207 | 0.3162 | 0.2630 | 0.4167 | 0.5075 |
| 10 | 0.4110 | 0.6277 | — | — | 0.7041 |
| 11 | 0.1604 | — | — | — | 0.5345 |
| 12 | 0.0277 | 0.1215 | 0.3206 | 0.4937 | — |
| 13 | 0.2497 | 0.2171 | 0.6439 | — | 0.8438 |
| 14 | nco | nco | 0.1953 | nco | 0.1381 |
| 15 | 0.0719 | 0.1199 | 0.2166 | 0.4089 | 0.6591 |
| 16 | 0.1540 | — | — | — | 0.4651 |
| 17 | 0.0357 | 0.0633 | 0.1971 | — | — |

The cases marked '—' were not considered due to very few observations for that $(k_1, k_2)$ pair. If there is a cross-over we estimate the proportion of decisions in the direction of the social cue 'CC', i.e. $k_2 = 10$ by the area under the fitted density between 6 and 11. No cross over cases are marked as 'noc'.

Table S13: Cross-over area under the influence of conflicting cue ‘FF’

| sub \ $(k_1, k_2)$ | (6,1) | (7,1) | (8,1) | (9,1) | (10,1) |
|---|---|---|---|---|---|
| 1 | — | — | 0.4928 | 0.4982 | 0.3434 |
| 2 | — | — | — | 0.5388 | 0.3926 |
| 3 | — | — | — | — | 0.5247 |
| 4 | 0.4117 | — | — | — | 0.5073 |
| 5 | 0.5217 | 0.3410 | 0.1830 | — | — |
| 6 | 0.3266 | 0.2361 | nco | — | — |
| 7 | 0.4153 | 0.4058 | — | — | 0.1602 |
| 8 | 0.4310 | — | nco | 0.2538 | 0.1938 |
| 9 | 0.5610 | 0.5466 | 0.5863 | — | — |
| 10 | 0.5157 | — | — | 0.7328 | 0.5577 |
| 11 | 0.3344 | — | — | — | 0.1281 |
| 12 | 0.6305 | 0.5799 | 0.5918 | 0.3035 | 0.0596 |
| 13 | 0.8309 | 0.6385 | 0.2783 | — | 0.5520 |
| 14 | 0.7860 | — | — | — | — |
| 15 | 0.3965 | 0.2553 | 0.1781 | 0.3729 | nco |
| 16 | — | — | — | — | 0.2214 |
| 17 | 0.4541 | nco | nco | 0.0825 | nco |

The cases marked ‘—’ were not considered due to very few observations for that $(k_1, k_2)$ pair. If there is a cross-over we estimate the proportion of decisions in the direction of the social cue ‘FF’, i.e. $k_2 = 1$ by the area under the fitted density between 1 and 6. No cross over cases are marked as ‘noc’.

Table S14: Comparison of cross over areas under conflicting and concurring cues

| Conf−Conc | p-value |
|---|---|
| $(1, 10) - (1, 1)$ | 0.0000 |
| $(2, 10) - (2, 1)$ | 0.0004 |
| $(3, 10) - (3, 1)$ | 0.0039 |
| $(4, 10) - (4, 1)$ | 0.2344 |
| $(5, 10) - (5, 1)$ | 0.0005 |
| $(6, 1) - (6, 10)$ | 0.0017 |
| $(7, 1) - (7, 10)$ | 0.0078 |
| $(8, 1) - (8, 10)$ | 0.0156 |
| $(9, 1) - (9, 10)$ | 0.0078 |
| $(10, 1) - (10, 10)$ | 0.0005 |

p-values of the right-tailed paired Wilcoxon signed rank test for the null hypothesis that A-B has zero median, where A contains the cross-over areas across the subjects under conflicting cue and B contains the cross-over areas across the subjects under concurring cue for the same value of $k_1$ at 5% significance level. It is evident that the cross-over is significantly more for conflicting cues.

Table S15: Two-way anova table

| Source | SS | df | MS | F | Prob>F |
|---|---|---|---|---|---|
| Columns | 0.00006 | 1 | 0.00006 | 0.04 | 0.8414 |
| Rows | 0.01006 | 3 | 0.00335 | 2.1 | 0.1031 |
| Interaction | 0.000143 | 3 | 0.00014 | 0.09 | 0.9659 |
| Error | 0.20419 | 128 | 0.0016 | | |
| Total | 0.21475 | 135 | | | |

Columns represent face/car, rows represent the 4 conditions (no comment, FF, FC/CF, CC) with 17 repetitions each. The differences are all insignificant as evident from the table.

Table S16: Significant voxels from statistical non-parametric test for concurring < conflicting using sLORETA during 200-250 ms post stimulus onset.

| X(MNI) | Y(MNI) | Z(MNI) | VoxelValue | Brodmann area | Lobe | Structure |
|---|---|---|---|---|---|---|
| -40 | 55 | 0 | -2.65E+00 | 10 | Frontal Lobe | Middle Frontal Gyrus |
| -50 | 15 | 10 | -2.59E+00 | 44 | Frontal Lobe | Precentral Gyrus |
| -45 | 15 | 10 | -2.56E+00 | 44 | Frontal Lobe | Precentral Gyrus |
| -55 | 20 | 10 | -2.54E+00 | 45 | Frontal Lobe | Inferior Frontal Gyrus |
| -45 | 50 | 5 | -2.50E+00 | 10 | Frontal Lobe | Middle Frontal Gyrus |
| -40 | 50 | 5 | -2.50E+00 | 10 | Frontal Lobe | Inferior Frontal Gyrus |
| -55 | 15 | 10 | -2.50E+00 | 44 | Frontal Lobe | Precentral Gyrus |
| -45 | 50 | 0 | -2.47E+00 | 10 | Frontal Lobe | Inferior Frontal Gyrus |
| -50 | 20 | 10 | -2.46E+00 | 45 | Frontal Lobe | Inferior Frontal Gyrus |
| -40 | 55 | 5 | -2.44E+00 | 10 | Frontal Lobe | Inferior Frontal Gyrus |
| -40 | 45 | 0 | -2.44E+00 | 10 | Frontal Lobe | Sub-Gyral |
| -40 | 55 | -5 | -2.43E+00 | 10 | Frontal Lobe | Middle Frontal Gyrus |
| -45 | 10 | 10 | -2.41E+00 | 44 | Frontal Lobe | Precentral Gyrus |

Table S17: Significant voxels from statistical non-parametric test for concurring < conflicting using sLORETA during 250-300 ms post stimulus onset.

| X(MNI) | Y(MNI) | Z(MNI) | VoxelValue | Brodmann area | Lobe | Structure |
|---|---|---|---|---|---|---|
| -40 | 55 | 0 | -2.41E+00 | 10 | Frontal Lobe | Middle Frontal Gyrus |
| -50 | 15 | 10 | -2.30E+00 | 44 | Frontal Lobe | Precentral Gyrus |
| -40 | 45 | 0 | -2.26E+00 | 10 | Frontal Lobe | Sub-Gyral |
| -45 | 15 | 10 | -2.26E+00 | 44 | Frontal Lobe | Precentral Gyrus |
| -40 | 50 | 5 | -2.25E+00 | 10 | Frontal Lobe | Inferior Frontal Gyrus |
| -45 | 50 | 5 | -2.24E+00 | 10 | Frontal Lobe | Middle Frontal Gyrus |
| -45 | 50 | 0 | -2.23E+00 | 10 | Frontal Lobe | Inferior Frontal Gyrus |
| -55 | 15 | 10 | -2.22E+00 | 44 | Frontal Lobe | Precentral Gyrus |
| -55 | 20 | 10 | -2.21E+00 | 45 | Frontal Lobe | Inferior Frontal Gyrus |
| -45 | 10 | 10 | -2.17E+00 | 44 | Frontal Lobe | Precentral Gyrus |
